# Dietary Human HSP90 Promotes Longevity and Healthspan in *Caenorhabditis elegans* via a Multi-Branched Stress Signaling Pathway

**DOI:** 10.64898/2026.11.11.765805

**Authors:** Thi Luong Vu, Sungwoo Kim, Han Choe

## Abstract

The loss of proteostasis is one of the primary drivers of aging. Despite the important role of the molecular chaperone HSP90 in maintaining proteostasis, its systemic role in aging appears contradictory because its inhibitors can effectively extend lifespan across diverse species. This study showed that dietary HSP90 supplementation via *Escherichia coli* significantly extended lifespan and healthspan of *Caenorhabditis elegans* without fertility trade-offs. Exogenous HSP90 likely acts as an extracellular signaling ligand that engages downstream kinase networks to drive the activation of three major longevity-associated transcription factors: DAF-16, SKN-1, and HSF-1. In addition, we demonstrated the additive effect of combining dietary HSP90 supplementation with the senolytic HSP90 inhibitor tanespimycin, indicating that extracellular proteostatic signaling and the intracellular inhibition of HSP90 can work together to enhance longevity. Finally, we engineered an antibiotic-free *E. coli* Nissle 1917 probiotic that constitutively expresses HSP90, which successfully extended the lifespan of *C. elegans*.

**Highlights:**

- Dietary delivery of human HSP90 extends *C. elegans* lifespan and healthspan.
- The longevity effect of exogenous HSP90 is driven by DAF-16, SKN-1, HSF-1, and mTORC2 signaling.
- Exogenous HSP90 and HSP90 inhibitors increase longevity additively.
- A stable, antibiotic-free probiotic was engineered for continuous HSP90 gut delivery.

## Introduction

As a result of a worldwide demographic shift, the total global number of older adults is expected to reach 1.4 billion by 2030 and 2.1 billion by 2050 ^1^. Effective therapies for this aging population must be developed in order to improve individual healthspan as well as reduce the burden on global healthcare systems. To achieve this, targeting the foundational hallmarks of aging is critical, and the loss of proteostasis is a notable primary driver ^2^. Disruption of proteostasis is marked by a significant impairment of protein folding, maintenance, and degradation, which leads to toxic protein aggregation and accelerated aging ^3,4^. To combat this catastrophic failure, molecular chaperone networks provide the constant surveillance required to maintain protein homeostasis by assisting *de novo* folding ^5,6^, refolding stress-damaged proteins ^6,7^, and ensuring strict quality control by targeting misfolded proteins for degradation ^8,9^.

Within this chaperone network, the HSP70 and HSP90 families are the most abundant under basal conditions ^10,11^. Therefore, targeting these two chaperones represents a promising therapeutic strategy to prevent the collapse of protein homeostasis during aging. Interestingly, previous research has demonstrated that the intranasal administration of exogenous recombinant human HSP70 to older mice successfully extends lifespan, rescues age-associated cognitive deficits, and mitigates cerebral lipofuscin accumulation ^12^.

The success of exogenous HSP70 provides a compelling proof-of-concept regarding the use of chaperones for aging prevention, establishing HSP90 as an equally promising therapeutic target. In contrast to the broad and non-specific manner in which HSP70 assists a large variety of protein folding processes ^13,14^, HSP90 functions in a highly specialized manner, serving as a molecular chaperone responsible for a distinct set of client proteins ^15,16^. HSP90 clients are predominantly signal transducers that are important in regulating metabolic homeostasis, stress responses, and cell proliferation processes ^11,17^. Therefore, the decline in HSP90 function leads to a systemic collapse of these regulatory networks ^11^, driving accelerated cellular senescence ^18^, immunosenescence, sarcopenia, and diverse pathologies rooted in the overloading of misfolded protein intermediates ^4,19^. Despite its essential role in cellular networks, the systemic administration of an HSP90 inhibitor unexpectedly extended healthy lifespan, delayed age-related pathologies, and downregulated the senescence marker p16^INK4a^ ^20^. The upregulation of HSP90 supports the anti-apoptotic pathways of senescent cells; therefore, the administration of an HSP90 inhibitor provides the benefit of clearing these senescent cells and results in reduced systemic inflammation ^20^. This leads to a conflict regarding whether to boost HSP90 function for organismal proteostasis or suppress it to achieve senolytic benefits.

To address this paradox, we used an inter-species dietary intervention to supply human HSP90 protein to *Caenorhabditis elegans*. Our results indicate that feeding worms with HSP90-expressing bacteria significantly extended both the lifespan and healthspan of the worms without compromising their reproductive fitness. Supported by our genetic epistasis analysis, we hypothesize that exogenous HSP90 may act as an extracellular signaling ligand that triggers the activation of endogenous DAF-16/FOXO, SKN-1/Nrf2, HSF-1, and the mTORC2 signaling hub. Importantly, this intervention has proven to be effective in overcoming the vulnerabilities of endogenous HSP90 (*daf-21*) mutants, signifying that our exogenous HSP90 supplementation is capable of compensating for the lack of endogenous chaperone function. Furthermore, when co-supplementing dietary human HSP90 with the HSP90 inhibitor, Tanespimycin (17-AAG), we observed an additive effect in lifespan extension, indicating that spatial compartmentalization, pairing intracellular senolytic clearance with extracellular proteostatic signaling, represents a complementary, rather than antagonistic, therapeutic strategy. Finally, in order to translate these findings into a viable clinical model, we successfully developed an antibiotic-free probiotic based on the *E. coli* Nissle 1917 strain to continuously deliver HSP90 in the uncontrolled gut, which extended the lifespan of *C. elegans*.

## Materials and Methods

### Materials

All chemicals and reagents utilized were of analytical or cell-culture grade. Carbenicillin disodium salt (as an alternative to ampicillin) and 1-thio-β-D-galactopyranoside (IPTG) were purchased from Carbosynth (UK). Streptomycin sulfate, kanamycin sulfate, sodium dodecyl sulfate (SDS), acrylamide/bis-acrylamide, and TEMED were acquired from Biosesang (Yongin, South Korea). Glycerol, ethanol, sodium hydroxide, yeast extract, dimethyl sulfoxide (DMSO), and guanidine hydrochloride were provided by Samchun Chemical (Pyeongtaek, South Korea). Sodium chloride, magnesium sulfate, magnesium chloride, and cholesterol were sourced from Sigma-Aldrich (St. Louis, MO, USA). Agar and sodium hypochlorite (NaOCl) were obtained from Junsei Chemical (Tokyo, Japan), while peptone and ammonium persulfate were supplied by Duksan Pure Chemicals (Ansan, South Korea). Coomassie Brilliant Blue R-250 and sodium azide were purchased from Amresco (Solon, OH, USA). Potassium dihydrogen phosphate and disodium hydrogen phosphate were obtained from Merck (Darmstadt, Germany). Additional reagents included Tris-base from Daejung Chemicals (Siheung, South Korea), agarose for gel electrophoresis from LPS Solution (Daejeon, South Korea), dithiothreitol (DTT) from Anaspec (Fremont, CA, USA), and 5-fluoro-2’-deoxyuridine (FUdR) from Biosynth (Staad, Switzerland). The HSP90 inhibitor Tanespimycin (17-AAG, Cat. No. S1141) was purchased from Selleckchem (Houston, TX, USA). All plasticware was sourced from SPL Life Sciences (Pocheon, South Korea).

The plasmid encoding the human heat shock complex (HSP90, HSP70, p60/Hop, Hsp40, and p23) was obtained from DNASU Plasmid Repository (DNASU ID: HsCD00813839) ^21^. The pET-28a(+) expression vector and the E. coli Origami 2 strain were sourced from Novagen (Darmstadt, Germany). The *E. coli* strain ClearColi BL21(DE3) was obtained from Lucigen (Middleton, WI, USA). The *E. coli* Nissle 1917 (EcN) strain expressing T7 RNA polymerase, EcN (T7) ^22^, was provided by DSMZ (DSM 115365). The *E. coli* strain (iVEC3) utilized for the *in vivo E. coli* cloning method ^23^ was provided by NBRP-*E. coli* at NIG.

### Construction of plasmids

To evaluate the individual effects of human heat shock proteins in C. elegans, each protein-coding sequence was subcloned into the pET-28a(+) vector backbone using the *in vivo Escherichia coli* cloning (iVEC) method ^23^. The gene fragment and the linearized vector were amplified by polymerase chain reaction (PCR). Both were designed with short homologous ends (20–30 bp) to ensure successful assembly via endogenous homologous recombination inside the bacterial cells. These fragments were then co-transformed into iVEC cells via electroporation, where the activity of the Exonuclease III (XthA) enzyme assembled them into complete plasmids. The transformed iVEC cells were then selected on agar plates containing kanamycin. The integrity of the expression vectors was verified by sequencing, conducted at Cosmogenetech (Seoul, South Korea).

To generate an antibiotic-free probiotic expressing HSP90, we used the overlap extension PCR (OE-PCR) method to join HSP90 fragments and a linearized vector containing the thyA-infA cassette at their homologous regions (30–40 bp). The assembled plasmids were then transformed into mutant *E. coli* DH10B Δ*thyA* and screened on solid M9t medium (1x M9 salts, 0.1 mM CaCl_2_, 2 mM MgSO_4_, 2% (w/v) glucose, 0.5% tryptone, and 0.01% (w/v) thiamine hydrochloride). Successful plasmid candidates were sequenced to verify the complete HSP90 open reading frame (ORF) at Cosmogenetech (Seoul, South Korea). To express HSP90 in the probiotic strain, the confirmed plasmid was transformed into *E. coli* Nissle 1917 Δ (*thyA, infA)* harboring the helper plasmid pR-essG. Transformants were incubated at 42°C in the presence of sucrose to cure the pR-essG plasmid via counterselection. The resulting strain successfully expressed HSP90 and was utilized for subsequent animal experiments.

### Protein Expression and SDS-PAGE Analysis

Plasmids were transformed into *E. coli* expression strains (ClearColi BL21(DE3), Nissle 1917 (T7), or Origami 2) via heat shock and selected on LB agar supplemented with either 50 µg/mL kanamycin or 100 µg/mL streptomycin. Single colonies were inoculated into LB broth containing the appropriate antibiotic and grown overnight at 37°C with shaking at 200 rpm. Cultures were diluted 1:50 into fresh LB broth (containing 50 µg/mL kanamycin or 100 µg/mL streptomycin) and incubated under identical conditions until reaching an OD600 of 0.6–0.8. Protein expression was induced with 1 mM IPTG, and cultures were further incubated at 30°C and 200 rpm for 6 hours. Uninduced control bacteria for lifespan assays were cultured in parallel. Both induced and control cells were harvested at the same final OD600.

Cells were harvested by centrifugation (13,500 × *g*, 1 min, 4°C) and lysed on ice via sonication (30% amplitude; 40 cycles of 1 s ON and 2 s OFF). Lysates were centrifuged at 13,500 × *g* for 10 min at 4°C to separate soluble and insoluble protein fractions. To evaluate protein expression and solubility, samples were mixed with 5X reducing sample buffer (250 mM Tris-HCl pH 6.8, 30% glycerol, 10% SDS, 0.01% bromophenol blue, 300 mM DTT) and denatured at 95°C for 20 min. Proteins were resolved using 10% Tricine SDS-PAGE and visualized by staining with Coomassie Brilliant Blue R-250.

### *C. elegans* maintenance

The wild-type *C. elegans* Bristol strain (N2) and all mutant or transgenic strains utilized in this study were obtained from the Caenorhabditis Genetics Center (CGC, University of Minnesota). A complete list of the strains and their corresponding genotypes is provided in Supplementary Table S1. *C. elegans* were grown on solid nematode growth medium (NGM) plates seeded with *Escherichia coli* OP50 as a standard food source, or specific bacteria for other experimental assays (e.g., feeding with engineered *E. coli* expressing HSP90). Routine stock cultures were maintained at 20°C; however, strains MQ130 [*clk-1(qm30)* III], CB1370 [*daf-2(e1370)* III], MQ887 [*isp-1(qm150)* IV], and JT6130 [*hsp-90(p673)* V] were maintained at 15°C to preserve stock viability.

### Age Synchronization and Lifespan Assay

The synchronization and lifespan assays were performed according to a previously described protocol ^24^, with some adjustments. Age-synchronized *C. elegans* populations were generated using an alkaline hypochlorite solution. Adult worms were recovered in 15 mL tubes by washing plates with M9 buffer, pelleted by centrifugation (400 × g, 2 min), and washed repeatedly with M9 buffer until the supernatant was clear. The worm pellet was then resuspended in 1 mL of M9 buffer and mixed with 1 mL of 2X alkaline hypochlorite solution (0.625 M NaOH, 1% NaOCl). The suspension was then shaken vigorously for 6 min, and the reaction was stopped with an excess volume of M9 buffer. After three additional washes with M9 buffer, the recovered eggs were incubated in 1 mL of M9 buffer at 20°C (or 15°C for the *daf-2(e1370)*, *clk-1(qm30)*, *isp-1(qm150)*, and *hsp-90(p673)* mutant strains) with gentle agitation for ∼24 h to allow for hatching and developmental arrest at the L1 larval stage.

Synchronized, arrested L1 larvae were plated onto NGM plates seeded with *E. coli* OP50 and grown at 20°C (or 15°C for the *daf-2(e1370)*, *clk-1(qm30)*, *isp-1(qm150)*, and *hsp-90(p673)* mutant strains). Upon reaching the late L4 stage, hermaphrodite worms were transferred to NGM assay plates supplemented with 49.5 µM 5-fluoro-2′-deoxyuridine (FUdR) to prevent progeny from hatching. These plates were pre-seeded with either induced *E. coli* expressing the target protein or uninduced control bacteria.

To prepare the food lawns for *C. elegans*, bacterial cultures were grown at 37°C and then induced with 1 mM IPTG (or left uninduced as a control) at an OD600 of 0.6–0.8. After adding IPTG, the bacterial cultures were continuously grown at 30°C to a final OD600 of ∼1.5. The bacterial cells were then pelleted at 6,000 × g for 15 min, washed twice with M9 buffer, and seeded onto the plates. In the case of the HSP90-EcN probiotic strain, the bacteria were cultured and harvested under the same conditions without the IPTG induction step.

Day 1 of the lifespan assay was designated as the day L4-stage larvae were transferred to FUdR plates. Standard lifespan assays were maintained at 20°C (including *hsp-90(p673)* mutants, which were shifted from 15°C to 20°C at the late L4 stage), except for the *daf-2*, *clk-1*, and *isp-1* strains, which were maintained at 15°C. *C. elegans* were transferred to fresh plates every 3 days and scored daily for viability. Death was defined as a lack of response to a gentle touch with a platinum wire. Worms that crawled off the agar, burrowed, or died from vulval rupture were censored from the survival analysis.

### Lifespan Assay of Combined Tanespimycin (17-AAG) and Exogenous HSP90

To evaluate the combined effects of dietary exogenous HSP90 supplementation and pharmacological HSP90 inhibition, lifespan assays were conducted using tanespimycin (17-AAG). This compound was solubilized in DMSO to a stock concentration of 10 mM, and right before plate pouring, the stock solution was added to the NGM medium to reach a final concentration of 25 μM. DMSO control plates were prepared using the same protocol with a final concentration of 0.25% DMSO to account for solvent toxicity. Both the 17-AAG and DMSO-control plates were supplemented with FUdR and seeded with either induced HSP90-expressing *E. coli* or the uninduced bacteria. Synchronization and lifespan assays were performed as previously described.

### Thrashing assay

Thrashing assays were conducted on Days 4, 7, and 10 of adulthood. On each testing day, nematodes (*n* ≥ 42) were placed individually in a 10 µL drop of M9 buffer on an agarose pad. Following a 1-minute adaptation period, their body bends were measured for 30 seconds under a Nikon microscope. Thrashing frequencies were subsequently analyzed using the wrMTrck plugin in ImageJ.

### Thermorecovery Survival Assay

Worms were synchronized and maintained as described in the lifespan assay section. On Day 7 of adulthood, plates containing the nematodes were subjected to a heat shock at 37°C for 2.5 hours, followed by a 24-hour recovery period at 20°C. Viability was scored, and death was defined as a lack of response to a gentle touch of platinum wire. Survival rates were calculated as the percentage of live worms relative to the total number of worms in each group.

### Fertility assay

Following synchronization, individual L4-stage hermaphrodites (*n* = 22 per condition) were transferred to individual non-FUdR NGM plates seeded with either induced HSP90-expressing *E. coli* or uninduced control bacteria. To accurately determine daily and total brood sizes, each adult worm was transferred to a fresh plate every 24 hours during its fertility window (five consecutive days). The plates containing the laid eggs were then incubated at 20°C for 48 hours to allow for hatching, and the hatched larvae were counted to determine the number of viable progeny.

### Quantification of Protein Aggregation

To evaluate protein aggregation levels, we used the transgenic *C. elegans* strain AM140 (rmIs132 [*unc-54p*::Q35::YFP]), which expresses Q35::YFP protein in the body wall muscles. These mutant worms were synchronized using an alkaline hypochlorite solution as described for the lifespan assays. When the worms reached the L4 stage, they were transferred to plates seeded with either HSP90-expressing E. coli or uninduced control bacteria, and were transferred to fresh plates every three days. At Day 4 of adulthood, animals were immobilized using 10 mM sodium azide on 2% agarose pads. Fluorescence images of the mutant worms were captured using a Celena X microscope (Logos Biosystems, South Korea) equipped with a 10× objective and a YFP filter. The total area of Q35::YFP aggregates and the total worm body area were measured using ImageJ software, and the aggregate area was divided by the corresponding worm body area to normalize for variations in animal size. A total of 81 worms per condition were analyzed across three independent biological replicates.

### Statistical analysis

The log-rank test was used to derive *P* values for lifespan assays via the Online Application for Survival Analysis 2 (OASIS 2) platform ^25^. Microsoft Excel was used to conduct two-tailed Student’s *t*-tests to compare two independent groups, and results are presented as the mean ± SEM. All experiments were conducted with at least three independent biological replicates, with the exact sample sizes (*n*) stated in the respective figure legends.

## Results

### Human HSP90 Multi-Chaperone Complex extended the lifespan of *C. elegans*

Misfolded and aggregated proteins accumulate with age due to the progressive decline in the function of the protein homeostasis network ^3^. Within this network, the HSP90 multi-chaperone complex is well known as the master regulator of proteostasis, governing numerous cellular processes ^16^. To investigate whether the HSP90 multi-chaperone complex could promote healthy aging in *C. elegans*, we transformed a multigene co-expression plasmid containing the core components of the complex (HSP40, p60/HOP, HSP90, p23, and HSP70; Fig. S1A)^21^ into the *Escherichia coli* ClearColi BL21(DE3) strain, and induced expression using 1 mM IPTG. Total cellular lysates and cell pellet fractions were analyzed via SDS-PAGE (Fig. S1B). The SDS-PAGE result confirmed the co-expression of all five chaperone components primarily in a soluble form, with only a minor insoluble fraction. Molecular weight bands were distinctly resolved, corresponding to each target protein: HSP90 (83 kDa), HSP70 (70 kDa), p60 (63 kDa), HSP40 (40 kDa), and p23 (19 kDa).

Following induction and the cell pellet collection, wild-type N2 *C. elegans* worms were fed ClearColi BL21(DE3) bacteria that expressed the complete HSP90 chaperone complex. The survival analysis of *C. elegans* revealed that the lifespan of those fed bacteria expressing the chaperone complex was significantly extended compared to those fed uninduced bacteria (Fig. 1A). The mean lifespan of the control worms was 17.3 days, whereas the mean lifespan of worms fed bacteria expressing the HSP90 chaperone complex was 22.4 days. These results indicated a statistically significant 29% increase in the mean lifespan of the worms (*P* < 0.0001), showing that dietary supplementation of the HSP90 complex could profoundly increase lifespan.

**Figure 1.**
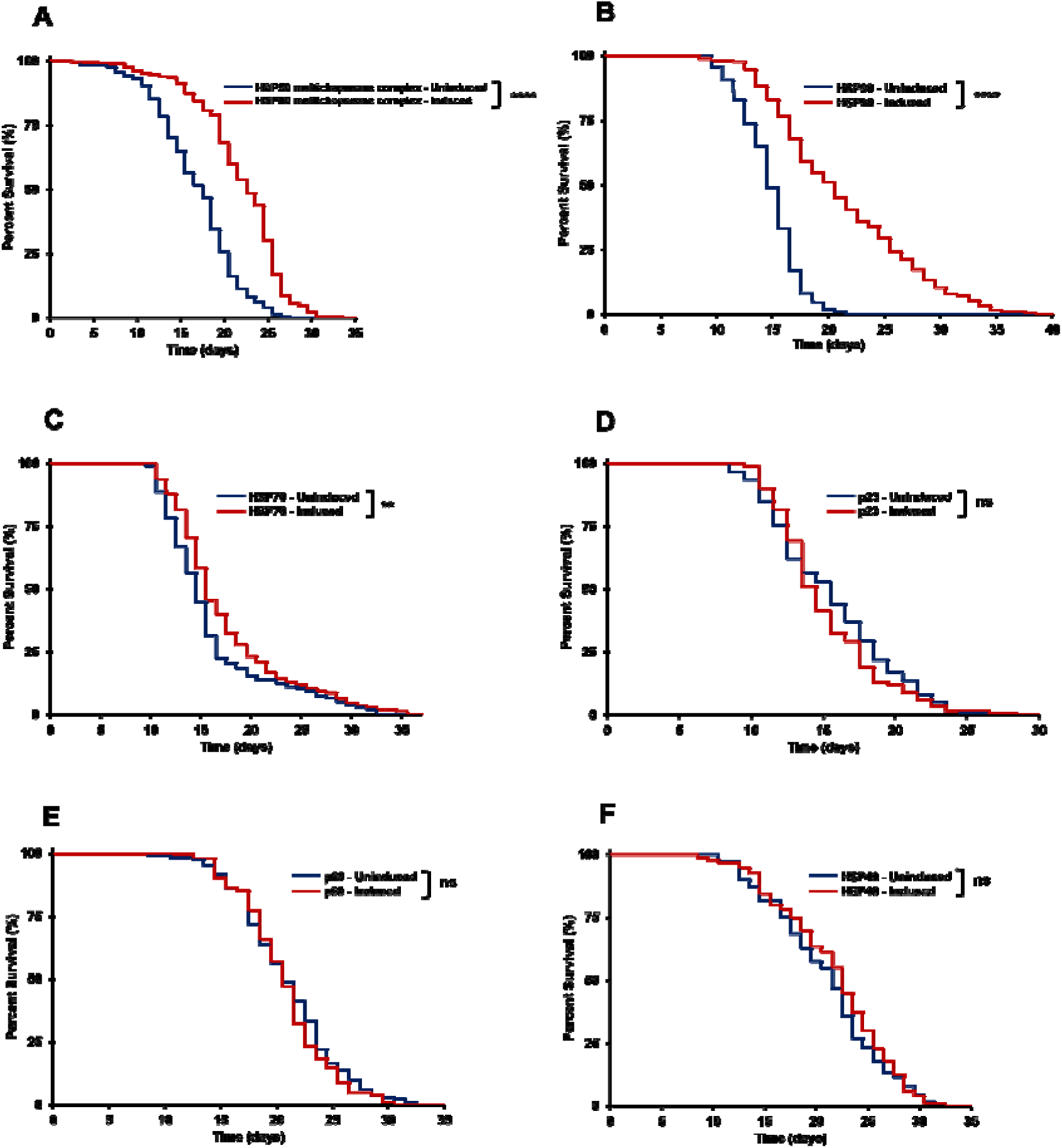
Lifespan extension of *C. elegans* is driven by a diet of bacteria expressing HSP90 and HSP70, but not HSP40, p23, or p60. Kaplan-Meier survival curves of wild-type (N2) worms fed *E. coli* ClearColi BL21(DE3) expressing specific human chaperone components. **(A)** The complete HSP90 multichaperone complex significantly extends lifespan (**** *P* < 0.0001; uninduced *n* = 191, induced *n* = 187). **(B, C)** Standalone expression of **(B)** HSP90 (**** *P* < 0.0001; uninduced *n* = 183, induced *n* = 172) and **(C)** HSP70 (\*\**P* < 0.01; uninduced *n* = 203, induced *n* = 214) extends lifespan. **(D–F)** Survival is not significantly altered (ns) in worms fed bacteria expressing the standalone co-chaperones **(D)** p23 (ns, *P* = 0.2956; uninduced *n* = 159, induced *n* = 166), **(E)** p60 (ns, *P* = 0.3068; uninduced *n* = 131, induced *n* = 102), or **(F)** HSP40 (ns, *P* = 0.3916; uninduced *n* = 111, induced *n* = 96). For all experiments, proteins were induced in the bacteria with 1 mM IPTG at 30°C (solid red lines) and compared to identical, uninduced control strains (solid blue lines). Statistical significance was calculated using the log-rank test. Data represent pooled results from three independent biological replicates.

### HSP90 as the Autonomous Driver of Longevity Within the HSP90 Chaperone Complex

Following the discovery of the lifespan extension provided by the complete HSP90 chaperone complex, further studies were conducted to identify which individual components were responsible for the survival benefit. Each of the protein sequences was isolated from the initial plasmid and subcloned into individual pET-28a(+) vectors using *in vivo E. coli* cloning (iVEC) (Fig. S2A). Upon individual induction in ClearColi BL21(DE3), we found that all proteins exhibited high levels of soluble expression (Fig. S2B) and were ready for use as a dietary source in *C. elegans* lifespan assays.

Lifespan assays for all five individual components of the human HSP90 complex (HSP40, p60/HOP, HSP90, p23, and HSP70) were performed. We found that bacteria expressing solely human HSP90 significantly increased the lifespan of *C. elegans* (Fig. 1B). The mean lifespan of worms fed HSP90-expressing *E. coli* was 21.7 days, compared to 15.2 days for control worms, representing a 43% extension in longevity (*P* < 0.0001). Dietary supplementation of human HSP70 also showed a reproducible but modest survival benefit in the lifespan of *C. elegans* (Fig. 1C). This increased the lifespan from 16.5 days in controls to 17.9 days in the test group, yielding an 8.8% extension in lifespan (*P* = 0.0065).

On the other hand, the remaining chaperone-expressing bacteria resulted in no extension in longevity. Worms fed p23-expressing bacteria exhibited lifespans statistically indistinguishable from controls (−2.0% change; mean lifespan of 15.7 days vs. 16.0 days; *P* = 0.2956). Similarly, dietary supplementation with p60-expressing bacteria (−1.35% change; 21.2 days vs. 21.5 days; *P* = 0.3068) and HSP40-expressing bacteria (3.5% difference; 22.2 days vs. 21.5 days; *P* = 0.3916) did not significantly alter survival (Fig. 1, D to F).

Because *C. elegans* lifespan is highly influenced by bacterial diet and growth conditions ^26–28^, we sought to confirm that these results were not strain-specific. The human HSP90 expression vector was transformed into varying bacterial backgrounds, including Origami 2 and an *E. coli* Nissle 1917 strain expressing T7 polymerase (EcN(T7)), and fed to worms. Exogenous HSP90 resulted in a reproducible lifespan increase across these distinct bacterial diets, yielding a 19% extension in the mean lifespan of *C. elegans* when fed HSP90-expressing Origami 2 (*P* < 0.0001) and a 14% extension when fed HSP90-expressing EcN(T7) (*P* < 0.0001; Fig. S3).

Collectively, these findings demonstrate that human HSP90 is the primary, autonomous driver of lifespan extension. Notably, its ability to extend lifespan without the presence of the co-chaperone proteins that are essential for protein folding processes supports the hypothesis that exogenous HSP90 functions independently of the traditional cellular chaperone pathway.

### Dietary Supplementation of HSP90 Improved Healthspan Without Reproductive Trade-offs

We also sought to determine whether HSP90-expressing bacteria improved physiological healthspan. We evaluated multiple aspects of healthspan, including neuromuscular function, stress resilience, systemic proteostasis, and reproductive profiles.

One of the most important aspects of aging is the profound decline in the neuromuscular system ^29,30^. We conducted a thrashing assay on Days 4, 7, and 10 of adulthood in both test and control groups to see if dietary supplementation with HSP90-expressing bacteria could mitigate age-related physical decline. In comparison to the control group (HSP90-uninduced), worms fed HSP90-expressing bacteria (HSP90-induced) exhibited a considerably higher bending rate (bends per 30 seconds) at all time points (*P* < 0.0001; Fig. 2A). Notably, this functional advantage became more pronounced with age. At Day 4, the bending rate of the control group was 52.1 ± 0.7, while that of the test group was 59.6 ± 0.7, representing an approximately 14% improvement in motility. However, this difference widened significantly at Day 7 to 27% (HSP90-uninduced = 41.1 ± 0.9; HSP90-induced = 52.2 ± 0.9). By Day 10 of adulthood, the bending rate of the control group had dropped to approximately 28 ± 1, whereas the worms fed HSP90-expressing bacteria maintained robust motility at 44 ± 1, representing a 57% improvement in motility. These results confirm that dietary HSP90 supplementation effectively mitigates age-related physical decline.

**Figure 2.**
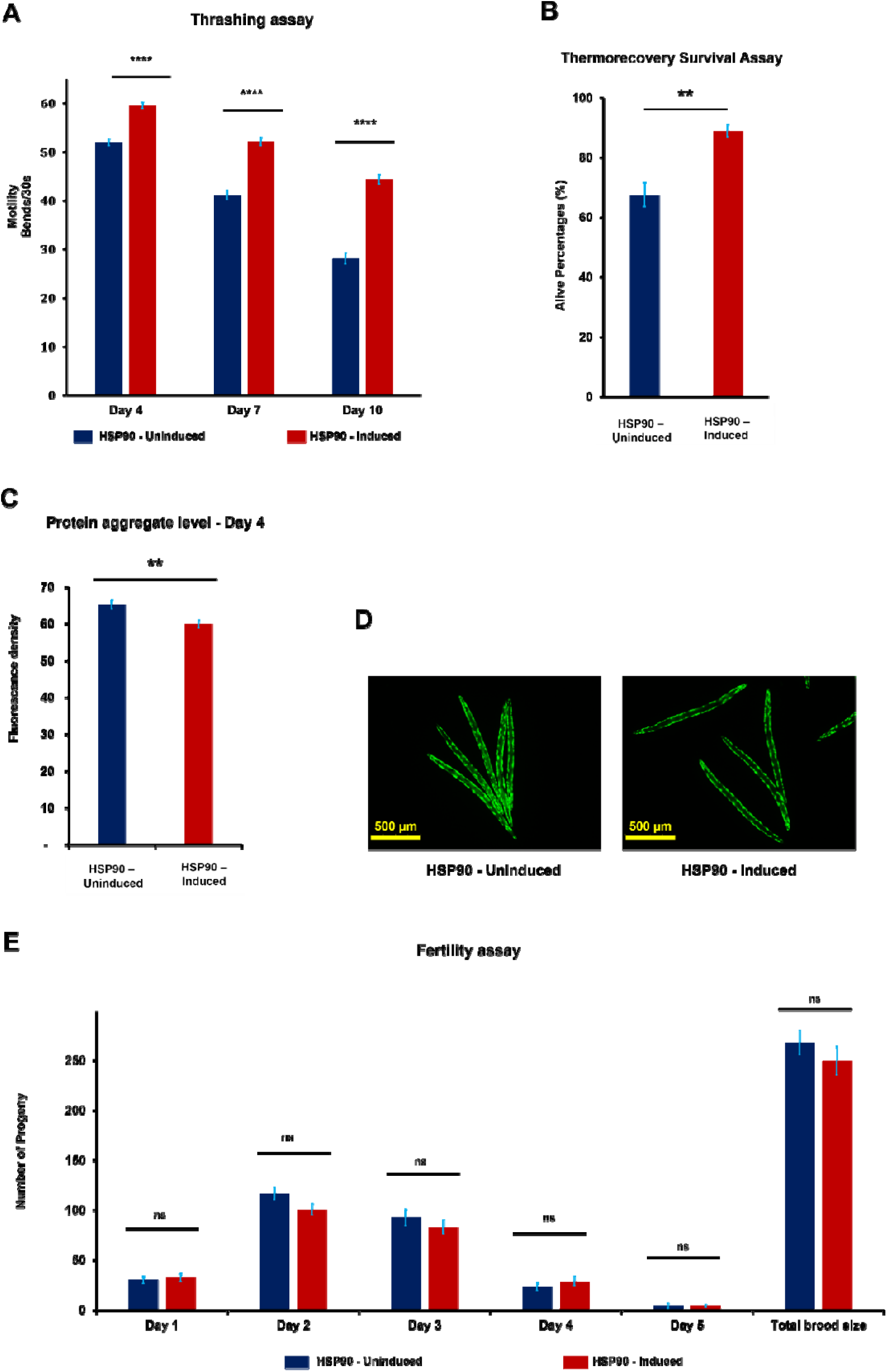
HSP90 expression in *E. coli* ClearColi BL21(DE3) improves motility, stress resistance, and protein homeostasis without altering fertility. In all assays, worms were fed *E. coli* ClearColi BL21(DE3) expressing HSP90 protein (induced with 1 mM IPTG) or an uninduced control. **(A)** Thrashing assay evaluating age-related motility at days 4, 7, and 10 of adulthood, quantified as body bends per 30 s (\*\*\*\**P* < 0.0001; Day 4: uninduced *n* = 71, induced *n* = 80; Day 7: uninduced *n* = 42, induced *n* = 52; Day 10: uninduced *n* = 83, induced *n* = 79). **(B)** Thermorecovery survival assay. Worms were subjected to heat stress (37°C for 2.5 h) followed by a 24 h recovery at 20°C. HSP90 induction significantly enhanced survival (\*\**P* = 0.00334; *n* = 85– 253 worms per condition; five independent biological replicates). **(C)** Quantification of Q35::YFP protein aggregation via relative fluorescence density at day 4 in AM140 (*rmIs132* [*unc-54p*::Q35::YFP]) animals, a model for age-progressive polyglutamine aggregation (\*\**P* = 0.00131; *n* = 81). **(D)** Representative fluorescence images corresponding to the quantification in panel (C), showing aggregates in day 4 AM140 worms (scale bar = 500 µm). **(E)** Fertility (brood size) assay showing daily progeny production (days 1–5) and cumulative total brood size. No significant differences were observed in daily reproductive timing or overall brood size (ns, *P* > 0.05 for all time points and total brood size; *n* = 22; four independent biological replicates). (For all panels, data are represented as mean ± SEM. Statistical significance was determined via Student’s *t*-test. Image analysis was performed using ImageJ).

In *C. elegans*, the ability to cope with severe environmental stress decreases as the proteostasis network progressively weakens with age ^31^. To assess whether dietary supplementation of HSP90 enhances stress resilience, we measured the survival rate of worms upon acute thermal shock. At Day 7 of adulthood, the worms were exposed to 37°C for 2.5 hours and then allowed to recover at 20°C for 24 hours. The overall survival rate was calculated by counting the live and dead nematodes. HSP90-expressing bacteria significantly enhanced the survival rate of the test group to 89% compared to 68% in the control group (*P* = 0.0033; Fig. 2B). These data suggest that supplementation with HSP90-expressing bacteria greatly improves the thermotolerance and systemic stress resistance of *C. elegans*.

In addition to an increased susceptibility to environmental stress, the loss of cellular proteostasis also results in the accumulation of toxic aggregated proteins in *C. elegans* ^31,32^. To determine whether HSP90-expressing bacteria could suppress protein aggregation, we used the AM140 (*rmIs132 [unc-54p::Q35::YFP]*) *C. elegans* strain, which expresses a polyglutamine (polyQ) protein fused to yellow fluorescent protein for visualization. Quantitative fluorescence imaging at Day 4 of adulthood demonstrated that the accumulation of cytotoxic polyglutamine foci in worms fed HSP90-expressing bacteria was significantly lower than in the control group (*P* = 0.0013; Fig. 2, C and D).

Extending lifespan often comes with reproductive costs ^33^; therefore, to test whether exogenous HSP90 supplementation induces a negative effect on fertility, we measured daily progeny production across a 5-day reproductive window. The HSP90-fed group and the control group exhibited no statistical difference in daily reproductive output (Day 1, *P* = 0.36; Day 2, *P* = 0.06; Day 3, *P* = 0.38; Day 4, *P* = 0.36; Day 5, *P* = 0.97) or total brood size (approximately 250-270 progeny per worm, *P* = 0.33; Fig. 2E). Collectively, our data indicate that the protective benefits of exogenous HSP90 supplementation do not significantly reduce or delay fertility.

Taken together, our results indicate that the dietary supplementation with HSP90-expressing bacteria extends not only *C. elegans* lifespan but also its overall healthspan. This intervention successfully preserved neuromuscular function, increased the worms’ ability to withstand acute environmental stress, and promoted the systemic clearance of age-related protein aggregates without any adverse reproductive trade-offs. In conclusion, exogenous HSP90 represents a promising and effective therapeutic strategy for delaying the aging process.

### Genetic Dissection of HSP90-Mediated Longevity Pathways

#### Exogenous HSP90 requires DAF-16 and HSF-1 for lifespan extension, acting independently of upstream IIS components and endogenous DAF-21

The insulin/IGF-1 signaling (IIS) pathway is highly conserved across diverse species, from simple invertebrates to mammals, with a profound impact on aging ^34^. In *C. elegans*, suppression of the DAF-2/IGFR drives the nuclear translocation of the DAF-16/FOXO transcription factor, thereby upregulating diverse stress-response and molecular chaperone networks ^35^. Given the central role of the IIS pathway in regulating aging, we investigated the relationship between exogenous HSP90-induced longevity and components of this pathway.

We first conducted the experiment on the *daf-2(e1370)* mutant strain, which carries a reduction-of-function mutation in DAF-2/IGFR and therefore increases the nuclear translocation of the DAF-16/FOXO transcription factor ^36^. Feeding HSP90-expressing bacteria to these mutant worms further extended their lifespan by 23%, from 29.6 days to 36.4 days (*P* < 0.0001; Fig. 3A). Similarly, dietary supplementation with HSP90 also promoted lifespan extension in the *age-1(hx546)* mutant worm strain. The *age-1* gene is a phosphatidylinositol 3-kinase (PI3K) homolog, and the AGE-1 protein functions directly downstream of the DAF-2 receptor in the IIS pathway ^37^; the *age-1(hx546)* mutant worm strain carries a reduction-of-function mutation in AGE-1/PI3K and lives longer than wild-type (N2) worms ^37,38^. HSP90-expressing bacteria still significantly extended the lifespan of this mutant strain, resulting in a 19% additive increase from 30.0 to 35.7 days (*P* < 0.0001; Fig. 3B).

**Figure 3.**
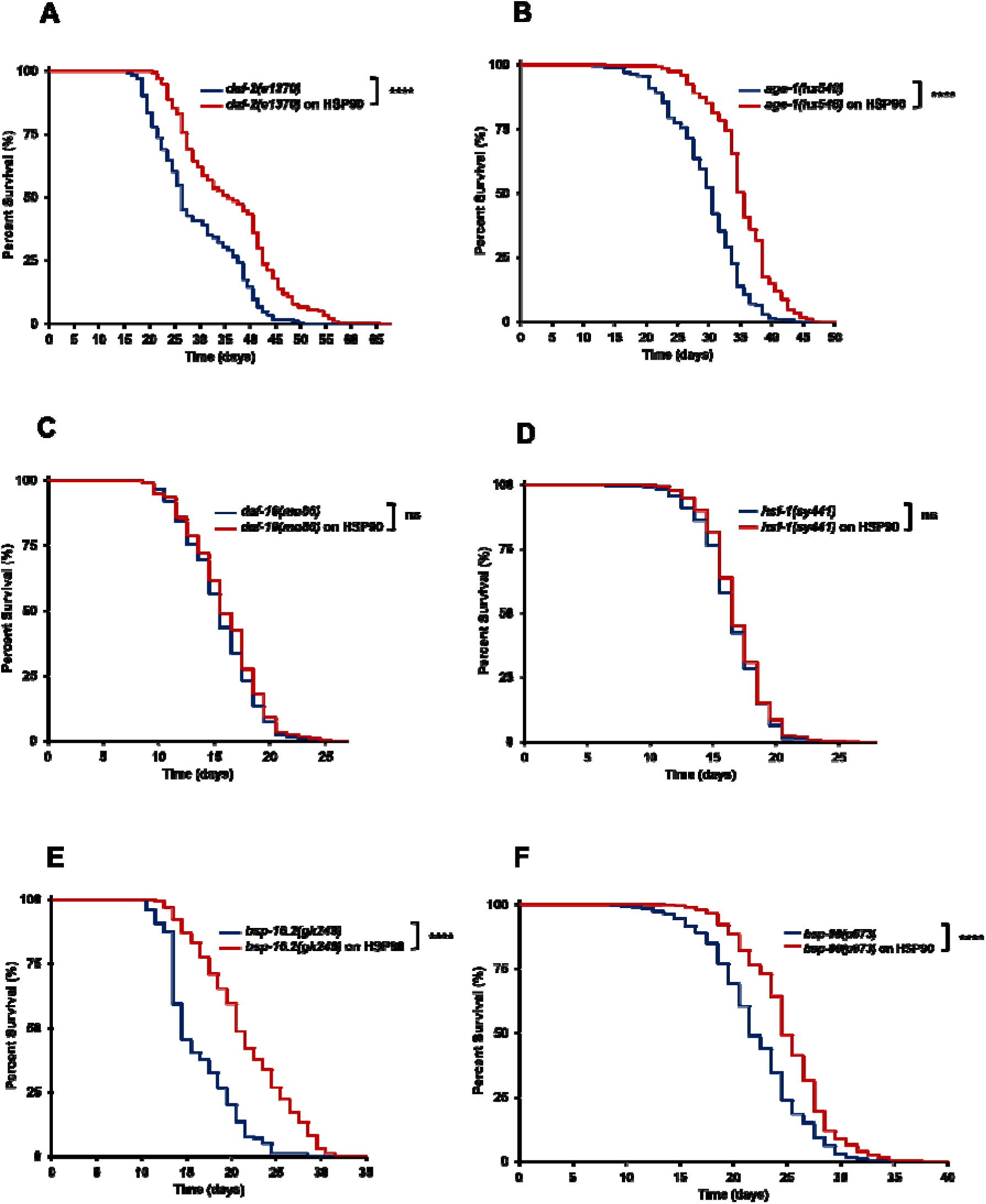
Dietary HSP90 extends lifespan independently of upstream IIS pathway but strictly requires DAF-16 and HSF-1. Dietary supplementation with exogenous human HSP90 extends the lifespan of *daf-2(e1370)*, *age-1(hx546)*, *hsp-16.2(gk249)*, and *hsp-90(p673)*, but not *daf-16(mu86)* and *hsf-1(sy441)*, mutant *C. elegans*. In each assay, worms were fed bacteria containing the human HSP90 expression plasmid, under either uninduced (blue lines) or induced (1 mM IPTG; red lines) conditions. **(A)** Kaplan-Meier survival curves demonstrate that HSP90 induction significantly extends the lifespan of *daf-2(e1370)* mutants maintained at 15°C (\*\*\*\**P* < 0.0001; uninduced *n* = 164, induced *n* = 177). **(B, E, F)** Similarly, HSP90 supplementation significantly extends the lifespan of **(B)** *age-1(hx546)* (\*\*\*\**P* < 0.0001; uninduced *n* = 308, induced *n* = 283) and **(E)** *hsp-16.2(gk249)* (\*\*\*\**P* < 0.0001; uninduced *n* = 153, induced *n* = 156), and **(F)** *hsp-90(p673)* (\*\*\*\**P* < 0.0001; uninduced *n* = 472, induced *n* = 570) mutants maintained at standard 20°C conditions. **(C, D)** In contrast, this survival benefit is abolished in **(C)** *daf-16(mu86)* null mutants (ns, *P* = 0.1018; uninduced *n* = 284, induced *n* = 289) and **(D)** *hsf-1(sy441)* mutants (ns, *P* = 0.0954; uninduced *n* = 615, induced *n* = 527) at 20°C. Statistical significance was calculated using the log-rank test. Data represent pooled results from three independent biological replicates.

Conversely, this lifespan-extending effect was abolished in the *daf-16(mu86)* mutant strain. Because the *daf-16* gene is knocked out, these animals lack functional DAF-16 protein and exhibit a significantly shorter lifespan compared to wild-type (N2) animals. Testing in this genetic background revealed that the survival curves of worms fed HSP90-expressing bacteria and the uninduced control group overlapped, showing only a 2% difference in mean lifespan (*P* = 0.1018; Fig. 3C). These results indicate that the lifespan extension provided by HSP90-expressing bacteria is dependent on the DAF-16 transcription factor, while its effect seems independent of upstream insulin/IGF-1 signaling components such as DAF-2 and AGE-1.

Previous studies have shown that HSF-1 and DAF-16 work synergistically to upregulate the expression of longevity-associated genes ^39–41^; on the other hand, endogenous DAF-21/HSP90 acts as a native repressor of HSF-1 under normal physiological conditions ^42,43^. Therefore, it is critical to determine how our exogenous HSP90 treatment interacts with this transcription factor. Lifespan assays using the *hsf-1(sy441)* reduction-of-function mutant strain demonstrated that dietary supplementation with exogenous HSP90 failed to provide any survival benefit. The mean lifespan of the group fed exogenous HSP90 (17.3 days) did not significantly differ from that of the control group (17.0 days, *P* = 0.0954; Fig. 3D) and the maximum lifespan was also similar (27 days vs. 26 days). The loss of benefit in *hsf-1(sy441)* mutant worms indicates that exogenous HSP90 is dependent on endogenous HSF-1 to extend lifespan.

Given this strict dependency on DAF-16 and HSF-1, we investigated whether the longevity effect of HSP90 relies on any single specific downstream effector protein activated by these transcription factors. We selected the *hsp-16.2(gk249)* null mutant strain, as HSP-16.2 is a canonical small heat shock protein and a classical biomarker of DAF-16 and HSF-1 ^39,44^. Feeding these mutant worms HSP90-expressing bacteria resulted in a 30% increase in mean lifespan compared to the control group (from 16.8 days to 21.9 days; *P* < 0.0001; Fig. 3E). This extension in lifespan indicates that while exogenous HSP90 requires the DAF-16 and HSF-1 transcription factors for its lifespan benefit, it is not strictly dependent on HSP-16.2, as the loss of this effector can likely be compensated by other functional cytoprotective proteins.

To determine whether exogenous human HSP90 functions independently or provokes the action of endogenous DAF-21/HSP90 for lifespan extension, we evaluated the survival of the *hsp-90(p673)* mutant strain fed HSP90-expressing bacteria. This strain is a *daf-21* reduction-of-function mutant, resulting in widespread physiological failures ^45^ and motility impairment ^46^. In contrast to the loss of survival benefit in *daf-16(mu86)* and *hsf-1(sy441)* mutants, HSP90-expressing bacteria increased the lifespan of *hsp-90(p673)* worms compared to the control group. The mean lifespan of the mutant worms fed HSP90-expressing bacteria increased by 12%, from 22.7 days in the uninduced control group to 25.5 days (*P* < 0.0001; Fig. 3F), and the maximum lifespan was extended from 35 to 39 days. This result demonstrates that exogenous HSP90 functions independently of endogenous DAF-21/HSP90.

#### The lifespan extension of exogenous HSP90 relies on intact mTORC2 signaling and the SKN-1 transcription factor

Beyond the insulin/IGF-1 signaling (IIS) pathway, we next investigated the interaction between dietary HSP90 and conventional metabolic nutrient-sensing signaling pathways. AMP-activated protein kinase (AMPK) is a metabolic regulator of cellular energy homeostasis. Serving as the primary cellular energy sensor, AMPK is activated by low energy levels and is essential for lifespan extension under caloric restriction ^47^. To evaluate the dependency of dietary HSP90 on the AMPK energy-sensing network, we used the *aak-2(ok524)* null mutant strain, a well-established model of AMPK deficiency. In the absence of functional AMPK, the lifespan-extending benefit of exogenous HSP90 remained intact; worms fed with bacteria expressing HSP90 had a mean lifespan of 18.1 days, which was 25% longer than that of the control at 14.5 days (*P* < 0.0001; Fig. 4A). These results indicate that exogenous HSP90 promotes longevity independently of the AMPK activation.

**Figure 4.**
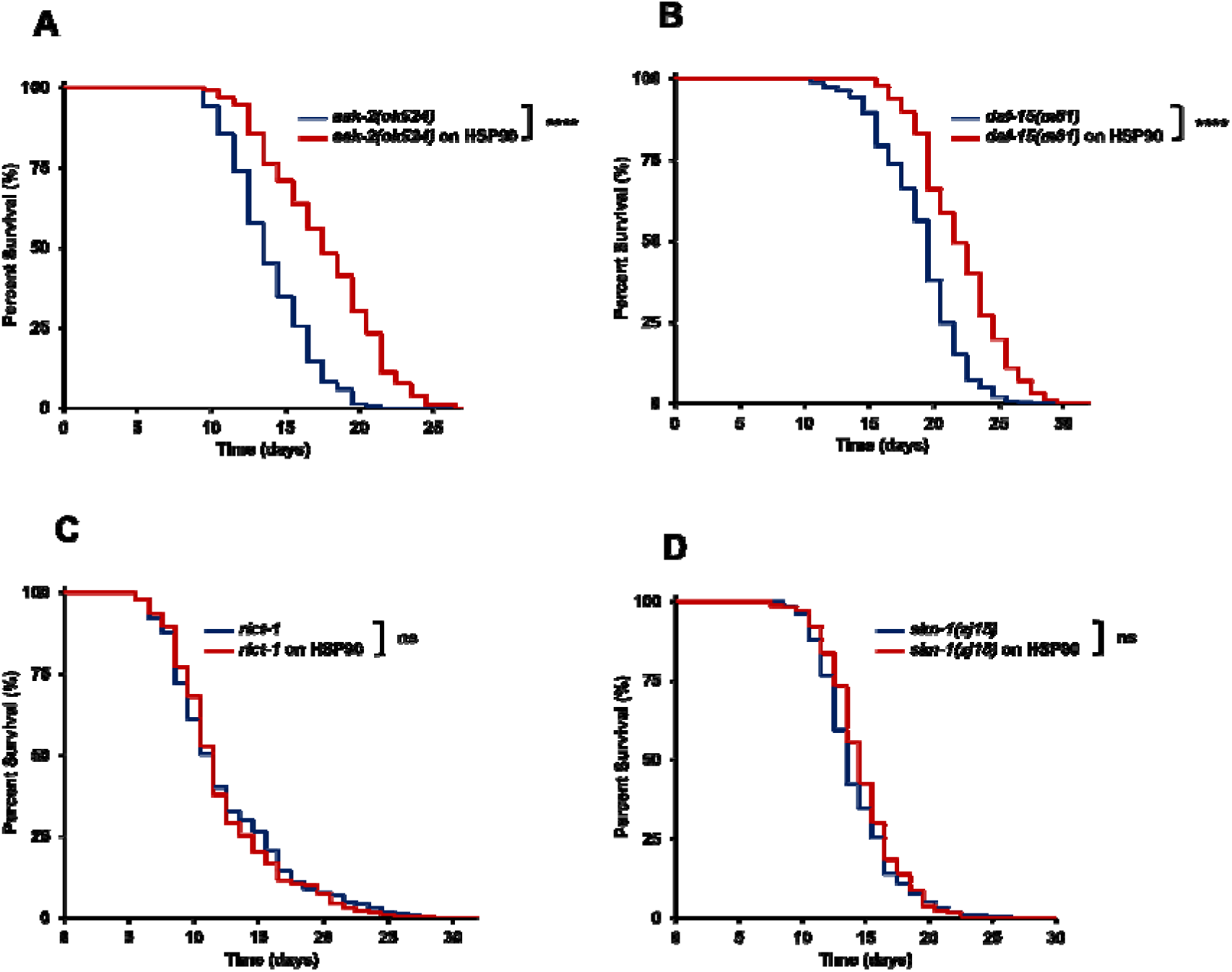
Dietary supplementation of human HSP90 extends *C. elegans* lifespan independently of *aak-2* and *daf-15* but strictly requires *rict-1* and *skn-1*. Dietary supplementation with exogenous human HSP90 extends the lifespan of *aak-2(ok524)* and *daf-15(m81)*, but not *rict-1(mg360)* or *skn-1(zj15)*, mutant *C. elegans*. In each assay, worms were fed bacteria containing the human HSP90 expression plasmid, under either uninduced (blue lines) or induced (1 mM IPTG; red lines) conditions. **(A-B)** Kaplan-Meier survival curves demonstrate that HSP90 induction significantly extends the lifespan of both (A) *aak-2(ok524)* (\*\*\*\**P* < 0.0001; uninduced *n* = 242, induced *n* = 232) and (B) *daf-15(m81)* (\*\*\*\**P* < 0.0001; uninduced *n* = 268, induced *n* = 309) mutants. **(C-D)** In contrast, HSP90 supplementation fails to significantly alter the survival of (C) *rict-1(mg360)* (ns, *P* =0.5135; uninduced *n* = 406, induced *n* = 398) and (D) *skn-1(zj15)* (ns, *P* = 0.1147; uninduced *n* = 215, induced *n* = 252) mutants. All assays were conducted at standard 20°C conditions. Statistical significance was calculated using the log-rank test. Data represent pooled results from three independent biological replicates.

The mechanistic target of rapamycin (mTOR) is an evolutionarily conserved signaling network that regulates cellular metabolism and growth in response to environmental nutrients and growth factors via two distinct complexes: mTORC1 and mTORC2 ^48^. High mTORC1 activity accelerates aging while eliminating its activation through caloric restriction, rapamycin, or genetic reduction can extend the lifespan significantly ^49^. On the other hand, mTORC2 is primarily regulated by growth factors rather than nutrient availability; since it serves as an upstream activator of several key survival kinases, inhibiting mTORC2 often results in a shortened lifespan ^50,51^. To investigate the relationship between exogenous HSP90 and mTORC1, we conducted a lifespan experiment using a worm strain with reduced mTORC1 activity (*daf-15(m81)unc-24(e138) IV*). In this mutant worm, exogenous HSP90 maintained its lifespan-extending benefit, resulting in a 16% lifespan extension (from 19.5 to 22.5 days; *P* < 0.0001; Fig. 4B). While exogenous HSP90 works independently of mTORC1, dietary supplementation of HSP90 failed to have a lifespan benefit in the mTORC2 mutant strain. In the lifespan experiment with *rict-1(mg360)* mutants (a model for mTORC2 deficiency with a loss-of-function in the defining RICT-1 subunit), the survival curve of worms fed HSP90-expressing bacteria was indistinguishable from the control, showing an insignificant change of −1.17% (*P* = 0.5135; Fig. 4C). These findings indicate that while HSP90-mediated longevity acts independently of mTORC1, its longevity benefits strictly require intact mTORC2 signaling.

Finally, to explore whether the lifespan-extending effect of exogenous HSP90 depends on the oxidative stress response, we conducted a lifespan experiment using the *skn-1(zj15)* mutant strain, which has significantly reduced functional SKN-1 protein. SKN-1 is a master transcriptional regulator of stress resistance and longevity whose nuclear translocation is strongly associated with cellular defense mechanisms. Loss-of-function SKN-1 mutants exhibit sensitivity to oxidative stress and significantly shortened lifespans ^52,53^. When studied in these mutant worms, exogenous HSP90 failed to extend lifespan; the mean lifespan of worms fed HSP90-expressing bacteria was 15.2 days, while the mean lifespan of the control group was 14.7 days, resulting in a non-significant difference of just 3.4% (*P* = 0.1147; Fig. 4D). Collectively, these results indicate that the lifespan extension induced by exogenous HSP90 is strictly dependent on mTORC2 and SKN-1.

#### Exogenous Human HSP90 Extends Lifespan Under Mild but Not Severe Mitochondrial Oxidative Stress

While low-grade mitochondrial stress promotes lifespan via beneficial hormetic signaling ^54^, severe dysfunction in mitochondria generates high levels of reactive oxygen species (ROS) that exceed cellular antioxidant capacity ^55^. To explore the effects of oxidative stress on the efficacy of exogenous human HSP90, we evaluated its impact on three electron transport chain (ETC) mutant strains.

First, we investigated the *clk-1(qm30)* mutant, a well-studied mitohormesis model that has slowed development and lives longer due to a deficiency in ubiquinone biosynthesis ^54,56^. Feeding the *clk-1(qm30)* mutant with bacteria expressing human HSP90 resulted in an additive longevity effect, with the mean lifespan increasing by 17% compared to the control group (from 26.2 to 30.7 days; *P* < 0.0001; Fig. 5A). This indicates that exogenous HSP90 signaling is independent of mitohormesis and functions efficiently even in the presence of mild endogenous oxidative stress.

**Figure 5.**
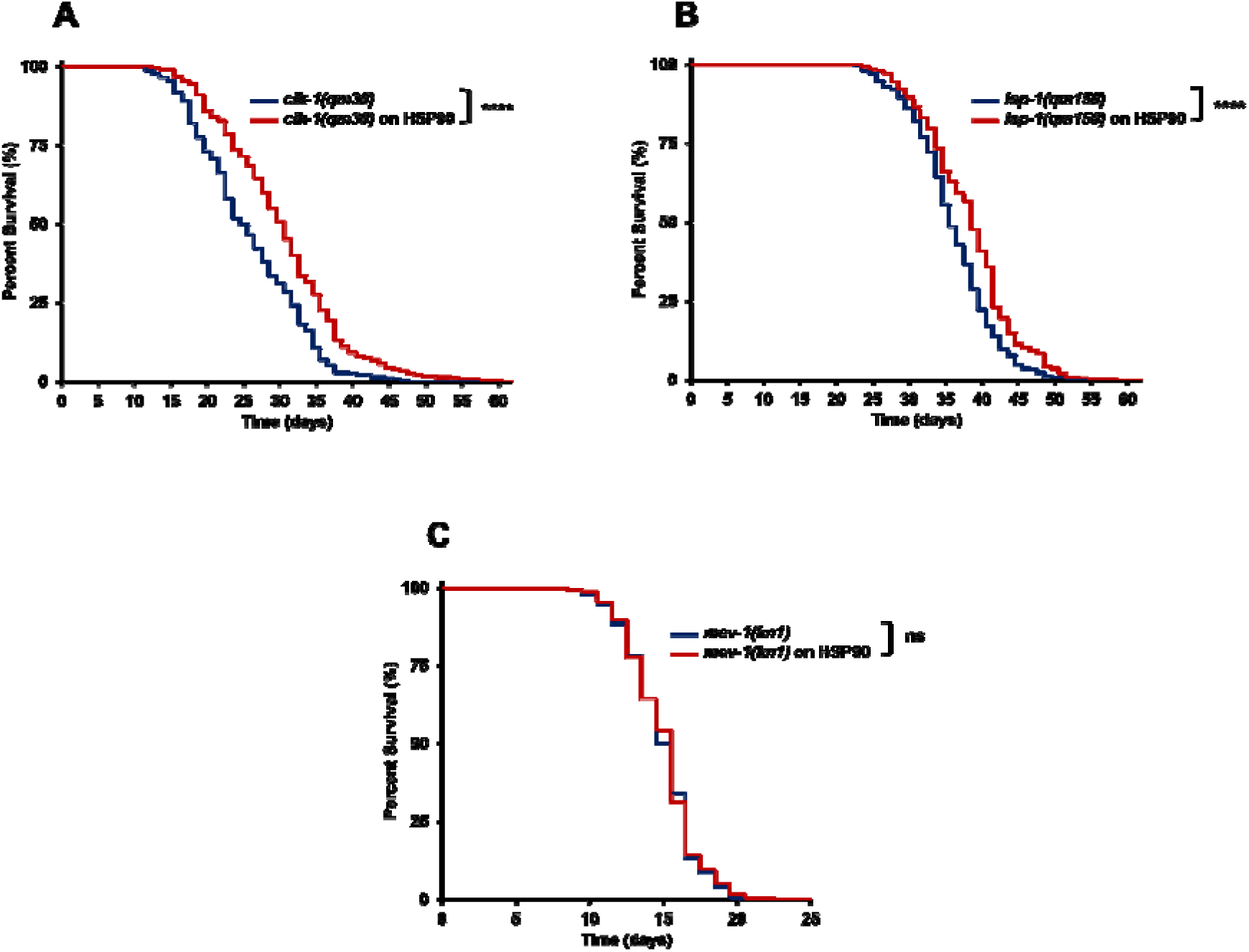
Exogenous human HSP90 extends *C. elegans* lifespan independently of *clk-1* and *isp-1* but fails to do so in *mev-1(kn1)* mutants. Dietary supplementation with exogenous human HSP90 extends the lifespan of *clk-1(qm30)* and *isp-1(qm150)*, but not *mev-1(kn1)*, mutant *C. elegans*. In each assay, worms were fed bacteria containing the human HSP90 expression plasmid, under either uninduced (blue lines) or induced (1 mM IPTG; red lines) conditions. **(A-B)** Kaplan-Meier survival curves demonstrate that HSP90 induction significantly extends the lifespan of both **(A)** *clk-1(qm30)* (\*\*\*\**P* < 0.0001; uninduced *n* = 268, induced *n* = 258) and **(B)** *isp-1(qm150)* (\*\*\*\**P* < 0.0001; uninduced *n* = 291, induced *n* = 341) mutants maintained at 15°C. **(C)** In contrast, HSP90 supplementation does not significantly alter the survival of *mev-1(kn1)* mutants (ns, *P* = 0.6897; uninduced *n* = 334, induced *n* = 309) maintained at standard 20°C conditions. Statistical significance was calculated using the Log-rank test. Data represent pooled results from three independent biological replicates.

We also evaluated another model of mitohormesis, the *isp-1(qm150)* mutant, which possesses a mutation in the Rieske iron-sulfur protein of Complex III ^57^. Similar to *clk-1*, the mutation in *isp-1* leads to ‘leakage’ in the electron transport chain, thereby reducing oxygen consumption and ATP production, which results in an increase in longevity driven by protective ROS signaling ^54^. Consistent with the *clk-1(qm30)* mutant, dietary supplementation with HSP90 significantly extended lifespan, increasing the mean lifespan by 6.3% (from 36.5 to 38.8 days; *P* < 0.0001; Fig. 5B).

Contrary to the positive additive effects observed in the mild mitohormetic models, this systemic protective effect was entirely lost in the *mev-1(kn1)* mutant background. The *mev-1(kn1)* mutant worms are defective in mitochondrial Complex II (succinate-coenzyme Q oxidoreductase), which causes mitochondrial abnormalities with continuous electron leakage, creating high levels of ROS and dramatically decreasing lifespan ^55^. Within this state of hyper-oxidative stress, feeding with HSP90-expressing bacteria failed to increase lifespan; the difference between the test group and control was not significant (0.5% increase; *P* = 0.6897; Fig. 5C). This suggests that the systemic cellular defense networks upregulated by exogenous human HSP90 are overwhelmed by severe, unmitigated endogenous oxidative damage.

Collectively, these data provide evidence for the limitations of exogenous HSP90 effects under different levels of mitochondrial stress. While HSP90 supplementation acted additively with the mild, protective ROS signaling in the *clk-1* and *isp-1* mutants to extend lifespan, it failed to protect against the harsh oxidative stress in the *mev-1* mutant. This indicates that while exogenous HSP90 functions independently of mitohormesis and safely bolsters survival alongside mild endogenous oxidative stress, its protective capacity cannot overcome the catastrophic production of ROS.

#### Additive lifespan extension by combining dietary exogenous HSP90 and endogenous HSP90 inhibition

A key paradox surrounding HSP90 research is that although its activity is important for protein homeostasis ^16^, the pharmacological inhibition of endogenous HSP90 also significantly extends lifespan, likely by selectively inducing cell death in senescent cells ^20^. To determine the interaction between exogenous HSP90 supplementation and HSP90 inhibition, we evaluated the combined effect of Tanespimycin (17-AAG) and dietary human HSP90.

The wild-type *C. elegans* N2 strain was maintained on plates supplemented with either 25 μM Tanespimycin or 0.25% DMSO as a control and fed either induced HSP90-expressing or uninduced control bacteria. Consistent with previous literature, Tanespimycin extended the lifespan by 11% compared to the Control group (from 17.7 to 19.6 days; *P* < 0.0001; Fig. 6A). Furthermore, aligned with the results we have previously shown, independent supplementation with exogenous HSP90 extended the mean lifespan of *C. elegans* from 17.7 to 22.8 days, showing a 29% increase over the uninduced bacterial control group (*P* < 0.0001; Fig. 6A). Taken together, these baseline experiments demonstrated that both pharmacological HSP90 inhibition and exogenous HSP90 supplementation independently extended lifespan in *C. elegans*, providing a strong foundation to evaluate their combined therapeutic potential.

**Figure 6.**
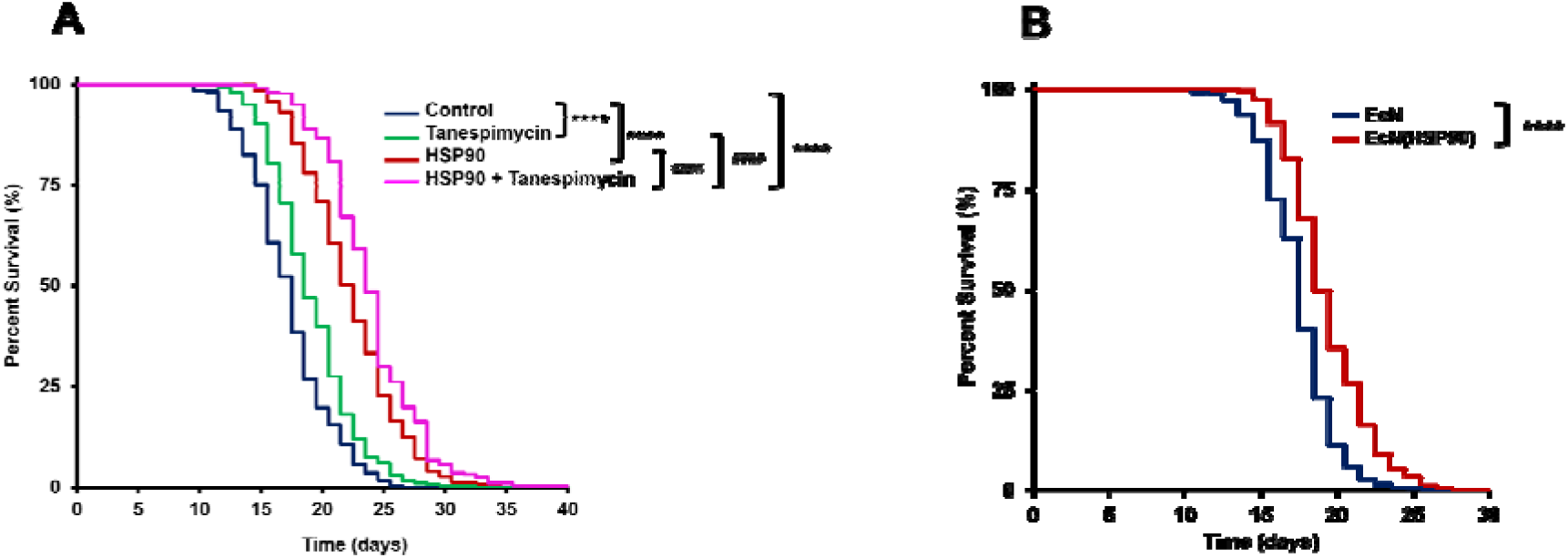
Additive lifespan extension by exogenous HSP90 and an HSP90 inhibitor, and the lifespan extension effect of an antibiotic-free HSP90-expressing EcN probiotic. **(A)** Survival curves of wild-type *C. elegans* treated with a combination of the HSP90 inhibitor Tanespimycin (25 µM) and dietary HSP90-expressing bacteria. The combined supplementation (*n* = 474) extended mean lifespan to 24.4 days, indicating an additive effect compared to both standalone Tanespimycin (*n* = 469, mean lifespan = 19.6 days) and standalone HSP90-expressing bacteria (*n* = 291, mean lifespan = 22.8 days). The vehicle control (DMSO + uninduced bacteria; *n* = 467) exhibited a mean lifespan of 17.7 days (\*\*\*\**P* < 0.0001 compared to the control group; ####*P* < 0.0001 for HSP90 + Tanespimycin compared to both standalone Tanespimycin and standalone HSP90 treatments). **(B)** Dietary supplementation with an antibiotic-free HSP90-expressing EcN probiotic significantly extended the lifespan of wild-type *C. elegans*. Lifespan assays demonstrated that HSP90-probiotic-fed worms (EcN(HSP90); *n* = 384) achieved a higher mean lifespan (20 days) compared to the control group carrying an empty vector (EcN; *n* = 413, mean lifespan = 18 days; ****P < 0.0001). Statistical significance was calculated using the Log-rank test. Data represent pooled results from three independent biological replicates.

Surprisingly, the combination of Tanespimycin with the human HSP90 diet resulted in an additive effect on lifespan. The worms treated with the combined approach exhibited the highest survival rate, with their mean lifespan increasing to 24.4 days, representing a 38% increase compared to the control group (17.7 days; *P* < 0.0001), a 24% increase compared to Tanespimycin alone (19.6 days; *P* < 0.0001), and a 7% increase compared to the standalone HSP90 diet (22.8 days; *P* < 0.0001; Fig. 6A). These results demonstrate that intracellular HSP90 inhibition and dietary exogenous HSP90 delivery do not antagonize each other; instead, they act additively to extend lifespan.

#### An antibiotic-free, HSP90-expressing probiotic extends *C. elegans* lifespan

To translate the healthspan and lifespan benefits of exogenous HSP90-expressing bacteria into a promising therapeutic, we developed an antibiotic-free probiotic that expresses HSP90. Standard laboratory bacteria depend on antibiotics to maintain their plasmids, making them highly impractical for uncontrolled environments such as the human gut ^58^. Moreover, this dependence presents an immense ecological and regulatory barrier to the development of live biotherapeutics due to the risk of horizontal gene transfer ^59,60^.

To create a safe and translatable HSP90 probiotic, we based our design on the antibiotic-resistance-gene-free plasmid (ARGFP) system ^61^ that utilized a double-knockout *Escherichia coli* Nissle 1917 (EcN Δ(*thyA*, *infA*)) strain. This mutant strain lacks two essential genes: the auxotrophic gene *thyA* and the essential translation initiation factor gene *infA*. Therefore, the survival of this strain heavily depends on an engineered plasmid containing both of these genes. By incorporating the HSP90 gene into the plasmid containing both essential genes, HSP90 protein can be stably expressed within bacterial cells in uncontrolled environments without the need for antibiotic selection. Furthermore, the HSP90 sequence was inserted under a constitutive promoter for constant expression without the need for chemical inducers. Ultimately, this approach allowed us to create a stable, antibiotic-free probiotic that continuously expresses HSP90 in the absence of selective pressure.

Consistent with our foundational experiments with the antibiotic-resistant *E. coli* strain, the HSP90-expressing EcN significantly extended the lifespan of wild-type *C. elegans* (N2) compared to the control group. The HSP90 probiotic increased the mean lifespan of the worms to 20 days compared to 18 days for the control, representing an 11% increase (*P* < 0.0001; Fig. 6B).

## Discussion

### The Central Role of HSP90 in Enhancing Systemic Proteostasis and Longevity

The maintenance of proteostasis is key to improving lifespan and healthspan, as it prevents protein aggregation during aging ^2,4,62^. Although systemic intranasal delivery of the protein chaperone HSP70 has been shown to extend the lifespan of mice ^12^, it remains unclear whether a similar treatment with the highly specialized chaperone HSP90 would also extend lifespan. On the one hand, HSP90 plays an essential role in regulating physiological processes, as its mutation or deficiency leads to abnormal development ^11,45,63^. Conversely, numerous studies have shown that treatment with HSP90 inhibitors can increase lifespan in certain models ^20,64^. This effect suggests that HSP90 might preserve deleterious cellular states and increase the risk of disease, thus masking its actual effects on lifespan.

Although feeding N2 worms the *E. coli* strain expressing the complete human HSP90 multichaperone complex (HSP40, p60 (HOP), HSP90, p23, and HSP70) increased lifespan by up to 29%, testing each component individually revealed significant differences in efficacy. Compared to the entire complex, dietary supplementation with human HSP90 alone resulted in a 43% lifespan extension. The difference in efficacy between the whole chaperone complex (29%) and HSP90 alone (43%) may be partly due to the metabolic burden of co-expressing five recombinant proteins, which could lower the yield of HSP90 delivered by the bacterial cells. On the other hand, dietary supplementation with exogenous human HSP70 still extended lifespan, but with a minor benefit of an 8.8% increase; our finding is consistent with previous observations of HSP70’s effects in mice ^12^. In contrast, HSP40, p23, and p60 failed to extend *C. elegans* lifespan. This lack of efficacy was unexpected given their critical roles: HSP40 is essential for baseline proteostasis ^65^ and its overexpression helps protect against lethality in *Drosophila* neurodegenerative models ^66,67^, while mutation of *sti-1* (the *C. elegans* homolog of p60) shortens lifespan ^68^ and the depletion of *sba-1* (p23) severely disrupts proteostasis ^65^. These results indicate that while HSP40, p23, and p60 are essential for intracellular protein folding, they lack the capability to directly drive longevity.

The lifespan-extending benefit of HSP90 was successfully replicated across *E. coli* strains, including ClearColi BL21(DE3), Origami 2, and *E. coli* Nissle 1917 containing a T7 promoter (EcN(T7)), demonstrating that this effect is driven by the HSP90 protein itself rather than by specific bacterial strains. Although the lifespan extension observed with ClearColi BL21(DE3) was greater than that observed with Origami 2 and EcN(T7), we hypothesize these variances stem from three main factors: endotoxicity, expression background, and intestinal packing and colonization dynamics. First is the difference in endotoxicity levels. Standard *E. coli* strains normally produce lipopolysaccharides (LPS) that contribute to chronic intestinal pathogenesis, while *ClearColi* is an endotoxin-free strain ^69^, providing a less toxic dietary baseline. Second, *ClearColi* is derived from the BL21(DE3) lineage, which lacks *lon* and *OmpT* proteases ^70^, facilitating the stable expression of recombinant proteins. In contrast, *Origami 2* is optimized for disulfide bond formation through specific mutations in the thioredoxin reductase (*trxB*) and glutathione reductase (*gor*) genes ^71^. Since HSP90 is a cytosolic chaperone protein that depends on conformational dynamics rather than disulfide bonds ^72^, the oxidizing environment of *Origami 2* may induce its misfolding or aggregation. Furthermore, it is highly likely that the yield of recombinant HSP90 varies across these distinct genetic backgrounds. Third, *ClearColi* exhibits a slower growth rate compared to conventional *E. coli* BL21(DE3), growing approximately 30% slower when cultured in defined media ^73^. This dynamic is particularly relevant for EcN(T7), which is modified from a wild-type probiotic that survives and proliferates more aggressively than laboratory *E. coli* strains ^74^. Such aggressive colonizations may cause physical stress and distension in the gut of *C. elegans*, thereby partially masking the lifespan-extending benefits of HSP90. Addressing these microbial dynamics, including the quantification of in vivo protein yield per strain, requires further investigation to successfully translate this approach into a promising therapeutic.

In conclusion, our results demonstrate that among the components of the HSP90 chaperone cycle, HSP90 itself is the primary driver of longevity, independent of specific *E. coli* strains. Furthermore, by using engineered HSP90-expressing bacteria for lifespan extension in *C. elegans*, we showed that dietary supplementation with the chaperone HSP90 alone was sufficient to extend lifespan, providing a promising therapeutic strategy for delaying aging.

### Exogenous HSP90 Improves Healthspan Without Reproductive Trade-offs

A common assumption in aging research is that increased lifespan correlates with an improved healthspan. However, many pharmacological and genetic anti-aging interventions extend lifespan but are accompanied by prolonged frailty ^75^. Our results across multiple physiological parameters demonstrate that exogenous HSP90 improves not only the lifespan but also the healthspan of *C. elegans*. Longitudinal tracking of motility throughout the adult lifespan revealed that while control worms showed a severe decline, dietary supplementation with HSP90 significantly mitigated this physical deterioration in advanced age. This demonstrates that exogenous HSP90 slows the rate of physical decline and actively preserves neuromuscular healthspan. Furthermore, HSP90-fed worms showed a strong capacity to maintain proteostasis, as evidenced by a significant clearance of cytotoxic polyglutamine foci *in vivo* within muscle tissues. Although dietary HSP90 is supplied via the gut, the efficient removal of aggregated proteins in distal muscle tissues suggests that exogenous HSP90 triggers a systemic protective response rather than acting locally within the intestine.

Previous studies have demonstrated that partial loss-of-function mutations in endogenous HSP90 (as seen in *daf-21(p673)* mutant worms) increase heat stress resistance ^42^, whereas tissue-specific overexpression of endogenous HSP90 makes the worms hypersensitive to heat stress ^76^. This dynamic is driven by a strict negative feedback loop: HSP90 functions as a negative regulator of HSF-1, and the overexpression of DAF-21/HSP90 results in HSF-1 inactivation, which damages the organism’s native stress response ^42,76,77^. In contrast to genetic *daf-21*-overexpressing mutant worms, our results showed that exogenous human HSP90 significantly enhanced acute heat stress resistance (89% survival vs. 68% in controls) in wild-type N2 worms. Indeed, exogenous HSP90 strictly requires HSF-1 to extend lifespan. We hypothesize that exogenous HSP90 likely functions as an extracellular ligand rather than an internalized chaperone. Therefore, it remains separated from cytosolic HSF-1 and may bypass this intracellular negative feedback loop. This suggests that instead of inhibiting HSF-1, as seen in *daf-21* overexpression mutants, dietary HSP90 acts as an extracellular signal in the gut, triggering a non-cell-autonomous cascade that systemically improves healthspan and thermal tolerance. Furthermore, our data show that exogenous HSP90 may activate DAF-16 and SKN-1, which are known to cooperate with HSF-1 to promote survival during heat stress ^39,78^. Finally, our acute 2.5-hour heat shock differs from previous 10-hour protocols ^76^; this variance in duration may also be one of the reasons for the contrasting outcomes.

Extending lifespan often comes with evolutionary trade-offs, such as delayed development or compromised reproductive ability, which are observed in previous long-lived genetic mutants or other life-extending interventions ^79,80^. However, with HSP90 supplementation, these healthspan improvements do not provoke reproductive trade-offs like those seen in *daf-2* mutant worms ^81^ or dietary restriction interventions ^82^. Our findings showed that even though there was a slight decrease in daily reproduction (Days 2 and 3) and in the total brood size of the group fed HSP90-expressing bacteria compared to the control, this overall decline was not statistically significant.

### Genetic Topology and Congruence with Endogenous Chaperone Pathways

To understand how exogenous HSP90 extends lifespan, it is valuable to review previous findings on its endogenous *C. elegans* homolog, *daf-21*. Because *daf-21* is essential for basic development, a loss of *daf-21* results in early larval lethality ^45^. On the other hand, a reduction-of-function mutation in *daf-21* results in a chemosensory deficiency caused by a failure in transmembrane guanylyl cyclase folding ^45^. Additionally, mutations in the *daf-21* gene impair reproduction due to its essential role in regulating the meiotic prophase/metaphase transition ^83^.

Beyond basic development, *daf-21* is heavily implicated in stress resistance and aging. DAF-21 has been proven to be essential for protecting *C. elegans* against stress, as it defends against environmental heavy metal toxicity ^84^ and promotes the nuclear translocation of transcription factor DAF-16 ^85^. Consequently, reducing DAF-21 function through genetic mutation or RNA interference reduces wild-type N2 lifespan and suppresses the longevity of long-lived mutants ^46,85,86^. Conversely, enhancing its activity through overexpression prevents proteostasis decline with age in *C. elegans* ^76^.

Our genetic dissection reveals that dietary supplementation with exogenous HSP90 likely activates endogenous protective pathways via specific transcriptional networks to drive longevity in *C. elegans*.

#### DAF-16 and HSF-1 are Required for the Pro-Longevity Effects of Exogenous HSP90

In long-lived IIS mutants, such as *daf-2* and *age-1,* endogenous DAF-21 and the chaperone network are essential downstream effectors for longevity ^85,86^. These studies suggest that HSP90 may function downstream of this signaling pathway. Our results revealed a similar logic: when introducing human HSP90 to upstream IIS mutants (*daf-2(e1370)* and *age-1(hx546)*), its pro-longevity effect was not abolished; instead, it provided an additive effect, driving a further lifespan extension in both strains. These results indicate that exogenous HSP90 may achieve its longevity benefits through a distinct mechanism that bypasses canonical IIS receptors.

Downstream of the IIS pathway, the transcription factors DAF-16 and HSF-1 act synergistically to drive the expression of longevity genes ^39,87,88^. In our study, the lifespan extension benefits of exogenous HSP90 were abolished in both the *daf-16(mu86)* and *hsf-1(sy441)* mutants. This strict dependence on both transcription factors for lifespan extension is very different from the behavior of endogenous HSP90/DAF-21. Endogenous cytosolic DAF-21 plays a dual, opposing role in cellular stress defense: while it is strictly required for DAF-16 localization ^85^, likely by stabilizing upstream stress-responsive kinases ^89^ such as JNK-1 ^90^ and CST-1 ^91^, it also functions as a potent HSF-1 repressor ^42,77^. Consequently, the intracellular overexpression of DAF-21 makes worms hypersensitive to heat stress ^76^. In contrast, our data demonstrate that exogenous human HSP90 protects worms from heat stress and likely promotes the nuclear translocation of both HSF-1 and DAF-16 to extend longevity.

This functional contrast led us to hypothesize that our dietary HSP90 supplementation may act as an extracellular signaling molecule. Indeed, HSP90 is well-documented in mammals as a secreted chaperone involved in cell migration and wound healing ^92–98^. Mammalian extracellular HSP90 (eHSP90) interacts with surface receptors such as LDL-receptor-related protein 1 (LRP1) and Toll-like receptor 2/4 (TLR2/4) ^96,99–101^ to trigger intracellular kinase cascades, including the proto-oncogene tyrosine-protein kinase, Src ^101,102^, and mitogen-activated protein kinase (MAPK) pathways ^103^. While HSP90 is well-documented as an extracellular signal in human and mouse models ^98,99,101^, we have no direct evidence yet that it acts as an extracellular signaling molecule in *C. elegans*. However, *C. elegans* possesses orthologs for known mammalian eHSP90 receptors, including LRP-1 (homologous to mammalian LRP1) ^104^ and the Toll-like receptor TOL-1 ^105^. Previous studies demonstrate that c-Jun N-terminal kinase (JNK) is a primary branch of the MAPK superfamily ^106^ known to physically interact with and enhance the nuclear translocation of DAF-16 ^107^. Regarding the HSF-1 transcription factor, other studies have identified p38 MAPKs, another branch of the MAPK superfamily, as highly efficient catalysts for the phosphorylation of HSF-1 at S326 ^108^. Therefore, we propose that our dietary HSP90 supplementation may act as an extracellular ligand that interacts with conserved surface receptors to stimulate downstream MAPK cascades, promoting the nuclear translocation of DAF-16 and HSF-1 independently of upstream IIS signaling.

Given the absolute requirement for DAF-16 and HSF-1, which are known to upregulate small heat shock proteins (sHSPs) to manage protein aggregation and extend lifespan ^39,86,87^, we next investigated the involvement of this downstream network using *hsp-16.2(gk249)* null mutant worms. In our study, exogenous human HSP90 extended lifespan independently of this canonical small heat shock protein, despite being absolutely dependent on the upstream DAF-16 and HSF-1 transcription factors. As a master transcriptional regulator, DAF-16 and HSF-1 co-orchestrate the expression of a large number of downstream stress-response genes, including antioxidants, autophagic effectors, and chaperones ^39,41,109^. The loss of a single downstream target, *hsp-16.2*, is unlikely to have a significant negative effect on this broad protective network. This proves that, as long as these primary transcription factors remain intact, the broad network of downstream protective genes is fully capable of compensating for the absence of *hsp-16.2*, securing proteostasis and driving longevity.

Finally, to determine whether exogenous human HSP90 simply provokes the endogenous nematode chaperone or acts independently, we evaluated the survival of the *hsp-90(p673)* mutant strain fed HSP90-expressing bacteria. This strain carries a reduction-of-function mutation in the endogenous *C. elegans hsp-90* (previously known as *daf-21*), resulting in sensory defects, decreased fertility ^45^, and impaired motility ^46^. Interestingly, dietary supplementation with human HSP90 significantly extended the lifespan of these mutant worms, demonstrating that the longevity benefit of exogenous HSP90 is independent of the endogenous chaperone. Furthermore, given its absolute requirement for the downstream DAF-16 and HSF-1 transcription factors, we hypothesize that exogenous HSP90 does not merely enter the cytosol to act as a replacement chaperone for protein folding; instead, it likely acts as an extracellular signaling ligand that triggers the nuclear localization of these transcription factors to promote longevity.

#### mTORC2 and SKN-1 are Required for the Pro-Longevity Effects of Exogenous HSP90

Beyond the IIS pathway, master metabolic kinases are crucial cellular sensors to determine cell fate in response to environmental factors, thereby profoundly impacting the aging process. Although intracellular HSP90 is essential for stabilizing and folding energy-sensing kinases, including AMPK and components of the mTORC1 complex ^110–112^, the longevity-promoting benefits of exogenous human HSP90 remain intact in both mutant strains that are deficient in mTORC1 (*daf-15(m81))* or AMPK signaling (*aak-2(ok524))*, proving that HSP90 functions independently of these critical metabolic checkpoints.

In contrast, when cultured at 20°C, the survival benefits conferred by exogenous HSP90 were abolished in *rict-1(mg360)* mutant worms, which carry a loss-of-function mutation in an essential subunit of mTORC2 ^51,113^. This demonstrates that mTORC2 is a strict epistatic block that HSP90 depends on for lifespan extension. Mechanistically, in mammalian models, the binding of extracellular Hsp90 to the LRP1 receptor results in increased phosphorylation of AKT at Ser473 ^114^. On the other hand, mTORC2 is the kinase responsible for phosphorylating AKT on this specific Ser473 residue ^115^. Together, this evidence strongly suggests that extracellular HSP90 triggers the activity of mTORC2. Furthermore, previous studies have demonstrated that in a cold environment (20°C), the cold-sensing TRPA-1 channel activates the mTORC2 in neurons, which subsequently phosphorylates serum/glucocorticoid regulated kinase 1 (SGK-1) to promote the nuclear localization of the DAF-16 transcription factor, thereby driving lifespan extension ^116,117^. Supported by the results of previous studies, we hypothesize that exogenous HSP90 acts as an extracellular signaling factor that mediates lifespan extension through an analogous pathway: eHSP90 → surface receptor → mTORC2 → SGK-1 → DAF-16 nuclear localization. This process may function dependently on, or independently of, TRPA-1 channel activity. This signaling pathway is consistent with earlier observations demonstrating that the life-extending effect of HSP90 is abolished in the *daf-16(mu86)* null mutant background. Taken together, these observations imply that exogenous human HSP90 does not rely on energy-sensing checkpoints such as AMPK and mTORC1, but strictly depends on the mTORC2–DAF-16 stress response pathway for longevity.

In parallel to this DAF-16 dependency, to determine whether exogenous HSP90 requires broader transcriptional regulators, we evaluated the Phase II antioxidant transcription factor SKN-1, which is responsible for the oxidative stress response ^52^. Strikingly, the pro-longevity effects of exogenous HSP90 were abolished in the *skn-1(zj15)* mutant strain. Previous studies have demonstrated that extracellular HSP90 localized to the cell surface results in MAPK activation ^103^. Notably, in *C. elegans*, PMK-1, a conserved member of the p38 MAPK family, acts as the upstream kinase responsible for phosphorylating and activating SKN-1 ^118^. Consequently, the lack of functional SKN-1 in the *skn-1(zj15)* mutant strain disrupts the proposed signaling cascade (eHSP90 → surface receptor → PMK-1 → SKN-1), thereby abolishing the survival benefits of exogenous HSP90.

In conclusion, we hypothesize that exogenous human HSP90 does not act as a general intracellular chaperone that repairs damaged proteins, but rather as an extracellular ligand that triggers receptor-mediated cellular stress responses. As an upstream extracellular signal, eHSP90 engages transmembrane cascades to control both the mTORC2–SGK-1 axis to activate the DAF-16 transcription factor, and the MAPK cascade to drive SKN-1-dependent antioxidant defenses. Consequently, the loss of functional mTORC2, DAF-16, or SKN-1 results in the loss of the HSP90-mediated survival benefit, firmly establishing this signaling network as the mechanism behind this longevity benefit.

#### Additive Survival Benefits and Limitations of Exogenous HSP90 Under Oxidative Stress

Mitochondrial damage results in the elevated production of reactive oxygen species (ROS), where different ROS levels dictate entirely different effects on lifespan. On one hand, severe ROS levels cause an accumulation of cellular damage; on the other hand, low-level ROS acts as a protective signal that can extend longevity (mitohormesis) ^119,120^. Previous studies have shown that endogenous cytosolic HSP90 interfaces with mitochondria via TOM70/TOM20 membrane receptors to mediate the import of proteins required for stress defense ^121,122^. Here, to understand the interaction between exogenous human HSP90 and mitochondrial respiration, we evaluated its pro-longevity effects across various electron transport chain (ETC) mutant worm strains. First, we used two mitohormetic models, the *clk-1(qm30)* and *isp-1(qm150)* mutants, in which the lifespan extension in these mutants is fundamentally driven by mild mitochondrial stress and metabolic deceleration ^54,57,123,124^. Our data showed that supplementing *clk-1(qm30)* and *isp-1(qm150)* mutants with human HSP90 resulted in additive longevity benefits. This additive effect suggests that exogenous HSP90 extends lifespan independently of mitochondrial stress regulation. This aligns with our previous hypothesis that exogenous HSP90 acts as an extracellular ligand and does not enter the cytosol to interact with mitochondrial TOM70/TOM20 receptors.

In contrast, the lifespan extension benefit of exogenous HSP90 was abolished when given to *mev-1(kn1)* mutants. The *mev-1(kn1)* mutant worms have compromised succinate dehydrogenase (ETC Complex II) activity, resulting in continuous electron leakage, elevated endogenous reactive oxygen species (ROS) levels, and hypersensitivity to oxidative stress ^55,125^. These results indicate that although exogenous HSP90 supplementation promotes longevity, it fails to rescue *mev-1(kn1)* mutant worms under hyper-oxidative conditions, demonstrating a clear physiological threshold. Taken together, although the protective benefits triggered by exogenous HSP90 were effective under moderate stress, they were insufficient to overcome the severe macromolecular damage and overwhelming oxidative stress driven by chronic Complex II failure.

#### Exogenous HSP90 Modulates Downstream Kinase Hubs to Drive the Concerted Activation of DAF-16, HSF-1, and SKN-1

Based on our collective genetic analysis, we propose that exogenous HSP90 extends lifespan through a multi-branched, integrated signaling network in *C. elegans* (Fig. 7). Upon ingestion, exogenous HSP90 bypasses the canonical DAF-2 insulin/IGF-1 receptor, a well-established regulator of longevity ^126,127^, to initiate other downstream protective signaling cascades. We identified three distinct kinase branches mediating this response. First, working in parallel with or dependent on the cold-sensing TRPA-1 channel in a 20°C environment, exogenous HSP90 may trigger the mTORC2 signaling hub in neurons. As established in previous studies, mTORC2 subsequently stimulates SGK-1 kinase to drive the nuclear localization of the DAF-16 transcription factor ^116,117,128^. Alongside mTORC2/SGK-1 signaling, we hypothesize that exogenous HSP90 triggers the JNK-1 signaling branch, which is known to further enhance the nuclear accumulation of DAF-16 ^90,129^. Parallel to the JNK-1/DAF-16 axis, exogenous HSP90 likely engages the PMK-1 (p38 MAPK) signaling branch to promote the concerted nuclear translocation of both HSF-1 ^108^ and the SKN-1 transcription factor ^130^. This is consistent with previous studies demonstrating that extracellular HSP90 interacts with cell surface receptors to trigger intracellular MAPK signaling, thereby driving SKN-1-dependent antioxidant defenses ^103^. Further investigations into these precise MAPK networks are needed to fully confirm this hypothesis. The concerted nuclear localization of DAF-16, HSF-1, and SKN-1 activates their downstream target genes to promote a cellular defense program that systematically extends healthspan and lifespan in *C. elegans* ^52,87^.

**Figure 7.**
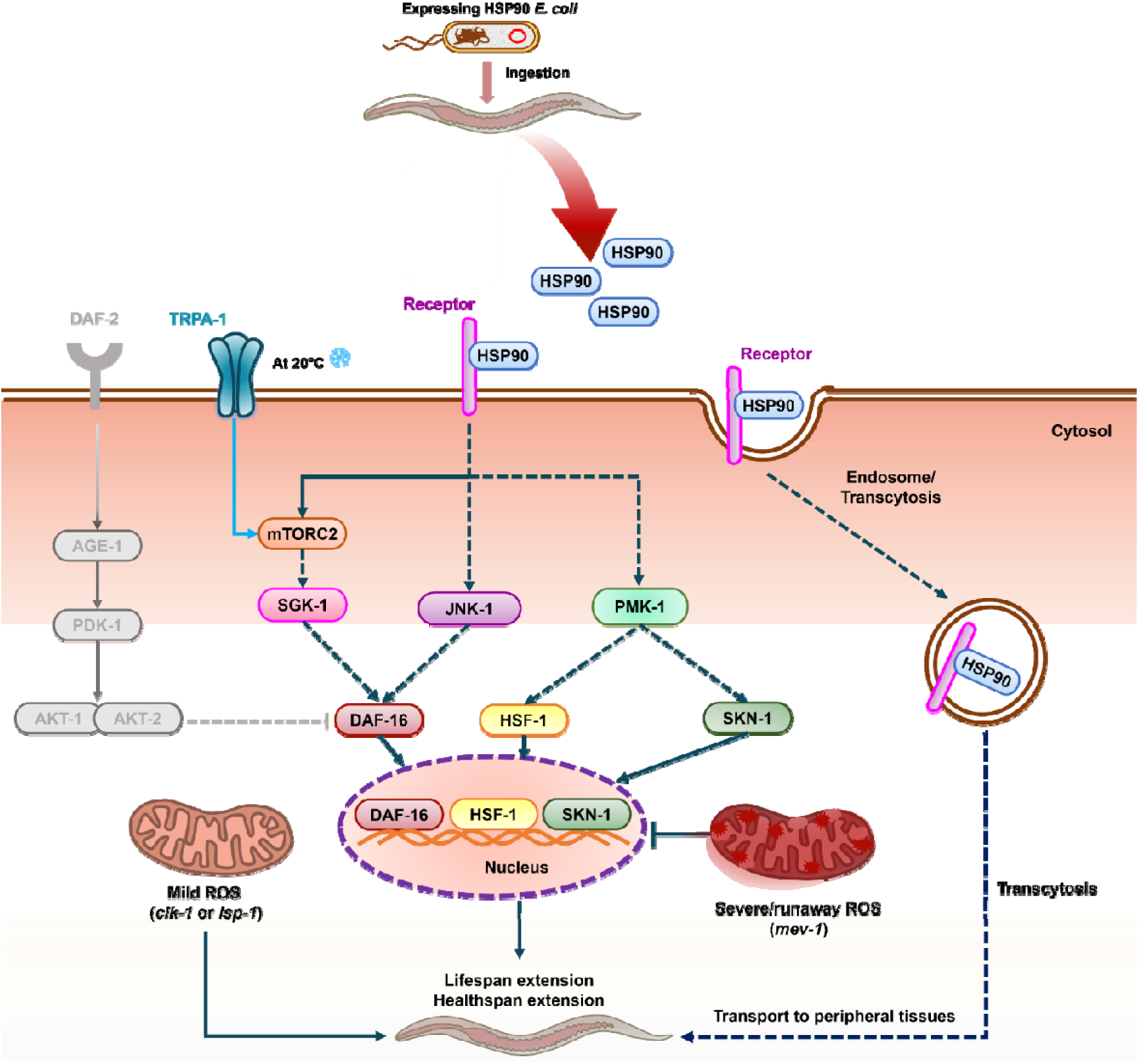
Proposed signaling mechanism for exogenous human HSP90-mediated lifespan and healthspan extension in *C. elegans*. Following the ingestion of *E. coli* expressing human HSP90, the exogenous chaperone is proposed to operate through a non-cell-autonomous mechanism that avoids the detrimental inhibition of endogenous HSF-1. Extracellular HSP90 may interact with an unidentified cell-surface receptor to activate intracellular kinase signaling, or alternatively, undergo endocytosis and transcytosis to peripheral tissues. This mechanism bypasses the canonical DAF-2 insulin/IGF-1 receptor cascade (grey) to drive lifespan and healthspan extension in *C. elegans* via three parallel signaling branches: ● **Pathway 1 (TRPA-1/mTORC2/SGK-1):** At 20°C, the TRPA-1 channel activates the mTORC2 complex and its downstream kinase SGK-1, thereby driving the nuclear translocation of the DAF-16 transcription factor. The presence of exogenous HSP90 may promote the maturation and proper assembly of mTORC2, resulting in the lifespan extension of *C. elegans*. ● **Pathway 2 (JNK-1/DAF-16):** Functioning in parallel, exogenous HSP90 is hypothesized to trigger the JNK-1 signaling branch, which acts as an intermediate regulator to further promote DAF-16 nuclear translocation, resulting in lifespan extension. ● **Pathway 3 (PMK-1/ HSF-1 & SKN-1):** Distinct from the JNK-1 regulatory network, exogenous HSP90 may promote the nuclear translocation of both HSF-1 and SKN-1 transcription factors, presumably through the canonical PMK-1 (p38 MAPK) signaling branch. Collectively, the synergistic nuclear localization of DAF-16, HSF-1, and SKN-1 induces the expression of stress-related genes to increase longevity and healthspan in *C. elegans*. This pathway functions in parallel and additively with mitohormesis but fails under severe oxidative stress. **Legend:** Solid arrows denote interactions proven through genetic epistasis experiments in this study (i.e., the dependence of the exogenous HSP90 longevity effect on mTORC2, SKN-1, DAF-16, and HSF-1). Dashed arrows denote putative receptors, transcytosis events, and unverified kinase intermediates (SGK-1, PMK-1, and JNK-1) based on established *C. elegans* literature.

From another perspective, the additive lifespan extension in *daf-2(e1370)* and *age-1(hx546)* mutants, combined with a strict requirement for a functional mTORC2 complex, suggests that exogenous HSP90 may bypass canonical upstream insulin/IGF-1 signaling to directly engage downstream kinase hubs. In mammalian models, eHSP90 interacts with surface receptors to recruit and activate Akt ^131,132^. Furthermore, Akt and SGK-1 belong to the same AGC kinase family ^133^ and share identical upstream regulators, including PDK1 ^134–136^ and mTORC2 ^115,137^. Therefore, we hypothesize that exogenous HSP90 likely triggers the mTORC2/SGK-1 axis in *C. elegans*. While SGK-1 normally inhibits DAF-16 ^138^ and SKN-1 ^139^, distinct receptor-mediated signals can reverse its function; instead of inhibiting, SGK-1 can drive DAF-16 nuclear translocation ^116,117^. A previous study also demonstrated that inhibiting AKT-1 activity by RNAi resulted in increased HSF-1 nuclear localization ^140^. Since AKT-1 and SGK-1 share similar consensus motifs ^141,142^, the functional flip of SGK-1 may not only promote DAF-16 translocation into the nucleus, but also indirectly promote the phosphorylation of DDL-1 and the dissociation of the DHIC complex, thereby promoting HSF-1 nuclear translocation. Once DAF-16 translocates to the nucleus, it directly upregulates the expression of *skn-1* ^143^ and works together with HSF-1 and SKN-1 to activate the expression of specific genes that promote longevity ^39^. Because the individual loss of DAF-16, HSF-1, or SKN-1 function abolishes the HSP90 survival benefit, the hypothesis that exogenous HSP90 targets a single downstream signaling cascade (such as the mTORC2/SGK-1 axis) elegantly explains this strictly codependent defense network without the need for parallel MAPK signaling.

Furthermore, based on our results in mitochondrial mutants, we found that the signaling network induced by exogenous HSP90 acts independently of pathways induced by mild ROS generation, allowing for additive survival benefits in mitohormetic models (*clk-1*, *isp-1*). However, this HSP90-mediated longevity faces a finite buffering capacity; it is ultimately overwhelmed by, and fails to protect against, the severe oxidative damage present in *mev-1* mutants.

#### Additive Action of Pharmacological Senolytics and Exogenous Human HSP90

HSP90 plays a central role in cellular protection and maintenance; it works as an ATP-dependent molecular chaperone that facilitates the folding of client proteins to ensure their stability and functionality ^16,144^. However, its influence extends far beyond basic protein maintenance. In *Drosophila*, mutations in *hsp90* disrupt homeotic gene expression ^145^, impair normal physical development, and lead to reduced fecundity and a shortened lifespan ^146^. In *C. elegans*, downregulation of *daf-21* (the homolog of *Hsp90*) causes defects in muscle structure ^46^ and fertility impairment ^45^, while a null mutation in *daf-21* results in early larval lethality ^45^. Conversely, while HSP90 plays an essential role in normal development and maintaining proteostasis, numerous studies have shown that its hyper-activated state promotes tumor growth by driving the survival, and invasion of cancer cells ^96,147,148^. Furthermore, the pharmacological inhibition of HSP90 has been shown to promote longevity in *C. elegans* ^149^, as well as extend healthspan and delay the onset of several age-related symptoms in mouse models ^20^. In mammals, HSP90 inhibitors function as senolytics by selectively inducing apoptosis in senescent cells, thereby mitigating their negative effects on the overall aging process ^20^. This conflict has created a biological tension in disease and aging research regarding whether upregulating or suppressing the HSP90 network is the optimal therapeutic strategy.

To navigate this paradox, we first evaluated the individual effects of these therapies on *C. elegans* lifespan. Notably, we found that exogenous HSP90 supplementation alone has a profound longevity effect compared to 17-AAG treatment alone. We hypothesize that several biological and physiological mechanisms account for this finding. By interacting with the N-terminal ATP-binding domain and blocking its ATPase activity, 17-AAG impairs HSP90 chaperone function and leads to the degradation of HSP90’s client proteins ^150,151^. Although this mechanism effectively induces the death of damaged, senescent and mutated cancer cells, it also poses a toxicity risk to normal tissues ^152,153^. Furthermore, lifespan experiments generally begin at the late L4 stage, when adult *C. elegans* consist of an invariant number of entirely post-mitotic somatic cells ^154,155^. Because HSP90 inhibitors rely on targeting hyper-activated HSP90 complexes typically found in highly proliferative or severely stressed cells ^148^, their therapeutic benefit in healthy, post-mitotic adult *C. elegans* is inherently limited. In contrast, our exogenous HSP90 likely acts as an extracellular signal that systematically upregulates the endogenous cellular pathway responsible for handling stress and maintaining proteostasis, safely supporting healthy post-mitotic tissues in *C. elegans* over their entire lifetime. Taken together, these findings suggest that exogenous HSP90 supplementation provides a safer and more effective strategy for maintaining proteostasis and extending lifespan in post-mitotic organisms compared to standalone HSP90 inhibitors.

Building on these individual effects, the combination of HSP90-expressing bacteria and the HSP90 inhibitor tanespimycin (17-AAG) provided an additive effect that extended lifespan further than either treatment alone. We hypothesize a dual-action mechanism to explain this synergy. In mammalian models, 17-AAG acts as a senolytic, targeting highly damaged, senescent cells, thereby alleviating systemic inflammatory and proteotoxic burdens ^20,64^. In post-mitotic organisms like *C. elegans*, which lack classical cellular senescence ^155^, 17-AAG extends lifespan primarily by disrupting the HSF-1/DAF-21 complex and thereby increasing HSF-1 transcriptional activity ^149,156^. In addition, the hyperactivation of DAF-21/HSP90 can stabilize toxic protein aggregates within aging tissues; 17-AAG mitigates this burden by inhibiting the activity of endogenous HSP90/DAF-21 ^149,157^. In parallel, the dietary delivery of exogenous human HSP90 likely acts as an extracellular signal to upregulate basal proteostatic networks across systemic tissues, thereby extending the healthspan and lifespan of *C. elegans*.

We propose that this synergistic dual-action is made possible by strict spatial and pharmacokinetic separation. 17-AAG is a small molecule that penetrates deep into somatic tissues to selectively inhibit hyper-activated endogenous HSP90 networks ^148,157^. Conversely, our dietary exogenous HSP90 is delivered via bacterial cells into the intestinal lumen, where it likely acts locally on the gut epithelium to initiate systemic, non-cell-autonomous stress signaling (e.g., the gut-to-neuron mTORC2 axis). This spatial compartmentalization allows 17-AAG to suppress internal toxic chaperone activity in peripheral tissues, whereas exogenous HSP90 safely acts as an external signaling ligand that promotes systemic cellular defense. Future *in vivo* studies are required to fully elucidate the precise tissue distribution and biochemical interactions driving this synergy.

Taken together, our data bridge the gap between HSP90 supplementation and HSP90 inhibition to offer a highly effective therapeutic approach. By pairing the clearance of intracellular proteotoxic burdens with systemic extracellular proteostatic support, this combined regimen offers a promising therapeutic strategy for targeting both aging and age-related diseases.

### Constitutive Expression of Human HSP90 in Antibiotic-Free Probiotics

Plasmids containing antibiotic resistance genes (ARGs) pose serious ecological and regulatory risks stemming from their uncontrolled dissemination via horizontal gene transfer ^158–160^. By successfully adapting the ARGFP system ^61^ to express HSP90 in *E. coli* Nissle 1917 (EcN) without the need for antibiotic selection, we eliminated the severe risks of ARG horizontal gene transfer. Within this antibiotic-free model, our data showed that the lifespan extension afforded by HSP90 in *C. elegans* was fully preserved. EcN has a well-established human safety record, offering proven therapeutic benefits for infectious diseases, metabolic disorders, and inflammatory bowel diseases (IBDs) ^161^. Therefore, utilizing this *E. coli* strain as a delivery chassis not only preserves the anti-aging potential of HSP90, but brings the inherent probiotic benefits of outcompeting pathogens and preserving the barrier integrity of the gut ^162,163^. From a bioengineering perspective, the lifespan extension driven by HSP90 in engineered EcN indicates that the metabolic cost involved in maintaining the dual-marker genes (*thyA*, *infA*) does not reduce the therapeutic payload capacity of the probiotic vector. This ARGFP system can efficiently synthesize and deliver human HSP90 to *C. elegans* without the need for chemical inducers or antibiotic maintenance. From a clinical and translational perspective, ARGFPs can be maintained in the mouse gut for at least 12 days ^61^, opening a promising proof-of-concept for safe, scalable chaperone delivery in animal models.

Taken together, we present the successful creation of an antibiotic-free live biotherapeutic that is highly scalable, safe, and feasible for sustained oral chaperone treatment, establishing a successful connection between fundamental proteostasis and microbiome engineering for preventing age-related decline.

### Mechanisms of Intestinal Transcytosis and Non-Cell-Autonomous Stress Signaling

Although we have observed the benefits of lifespan extension and healthspan improvements in *C. elegans* following dietary supplementation with HSP90, the precise mechanism by which the 83 kDa recombinant human HSP90 crosses the *C. elegans* intestinal lumen remains unknown. However, previous studies have shown that ingested proteins are not always entirely degraded by intestinal proteases; rather, they can undergo apical endocytosis and translocate from the gut lumen into systemic circulation ^164,165^. Importantly, this uptake mechanism is capable of transporting large macromolecules, such as fluorescently labeled proteins ^166^. Furthermore, therapeutic studies have successfully extended lifespan in mammalian models via the intranasal administration of exogenous recombinant human HSP70 chaperones ^12^. Thus, while intestinal transcytosis may enable the transfer of a fraction of human HSP90 across the gut barrier to access distal tissues ^167–169^, a highly plausible alternative is that exogenous HSP90 initiates non-cell-autonomous signaling directly from the gut surface.

Supporting the hypothesis of a cross-tissue effect of exogenous HSP90, our results demonstrate that the lifespan extension benefits of exogenous HSP90 require mTORC2 signaling at 20°C.

Previous research has confirmed that at lower temperatures (20°C), RICT-1/mTORC2 promotes the nuclear localization of DAF-16 in neuronal tissue, resulting in lifespan extension effects ^117^. Therefore, the reliance on mTORC2 suggests that exogenous HSP90 may trigger an inter-tissue endocrine or kinase cascade from the gut to neuronal tissue. Indeed, the intestine serves as the command center for systemic stress resistance in *C. elegans* ^170–172^. Consequently, even if exogenous human HSP90 activates MAPK and mTORC2 networks locally within the gut, it can trigger a non-cell-autonomous signaling cascade in distant peripheral tissues ^170^. This may provide an explanation for the alleviation of muscular aggregates, even if HSP90 chaperones remain in the gut ^76,173^.

## Conclusion

Loss of proteostasis is one of the primary drivers of aging, yet therapeutic targeting of the HSP90 chaperone remains controversial. Here, we demonstrated that dietary supplementation of human HSP90 significantly extended the lifespan and healthspan of *C. elegans* compared to other co-chaperones. We hypothesize that exogenous HSP90 functions as an extracellular signaling ligand that triggers intracellular signaling via the TRPA-1/mTORC2/SGK-1 kinase network to promote the nuclear translocation of DAF-16, SKN-1, and HSF-1 transcription factors. This receptor-mediated mechanism compensates for the age-related decline in proteostasis without inducing the thermal hypersensitivity seen in previous literature. However, the exact cell-surface receptor for exogenous HSP90 remains unidentified. While exogenous HSP90 supplementation extended lifespan additively with mitohormesis, it failed under severe oxidative stress. Interestingly, we observed an additive effect on lifespan extension when combining the intracellular HSP90 inhibitor 17-AAG with dietary supplementation of HSP90. Finally, we successfully engineered an antibiotic-free *E. coli* Nissle 1917 probiotic to continuously deliver human HSP90, establishing a translatable biotherapeutic for aging.

## Data Availability

All data supporting the findings of this study are available within the article and its Supplementary Information files. Additional raw data and materials (including engineered plasmids and bacterial strains) are available from the corresponding author upon reasonable request.

## Author Contributions

T.L.V. and H.C. conceived the project and designed the experiments. T.L.V. performed the plasmid construction, bacterial protein expression, and *C. elegans* lifespan and healthspan assays. T.L.V. analyzed the data, prepared the figures, and wrote the original draft of the manuscript. S.K. and H.C. critically reviewed and edited the manuscript. H.C. supervised the study and acquired funding. All authors have read and agreed to the published version of the manuscript.

## Supplementary

### Supplementary Figures

**Figure S1.**
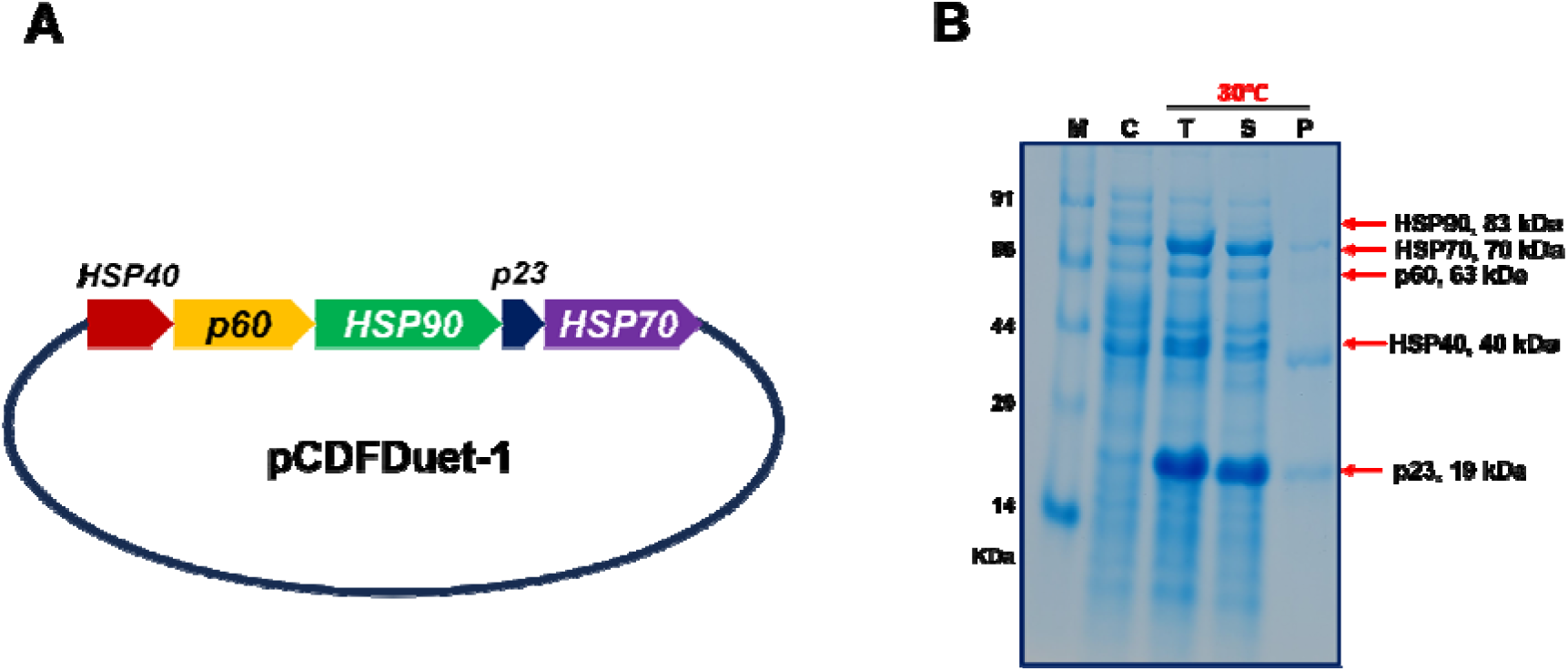
Construction and expression of the human HSP90 multichaperone complex in *E. coli* ClearColi BL21(DE3). **(A)** Schematic of the pCDFDuet-1 expression vector engineered to co-express the components of the human HSP90 multichaperone system (HSP40, p60, HSP90, p23, and HSP70). **(B)** Co-expression of the HSP90 multichaperone complex was analyzed by SDS-PAGE. Multichaperone expression was induced with 1 mM IPTG at 30°C; the control was the identical bacteria containing the vector but lacking IPTG induction. Lanes: M, molecular weight marker; C, uninduced control; T, total cell lysate; S, soluble fraction; P, insoluble pellet. Red arrows indicate the expected molecular weights of the individual components: HSP90 (83 kDa), HSP70 (70 kDa), p60 (63 kDa), HSP40 (40 kDa), and p23 (19 kDa).

**Figure S2.**
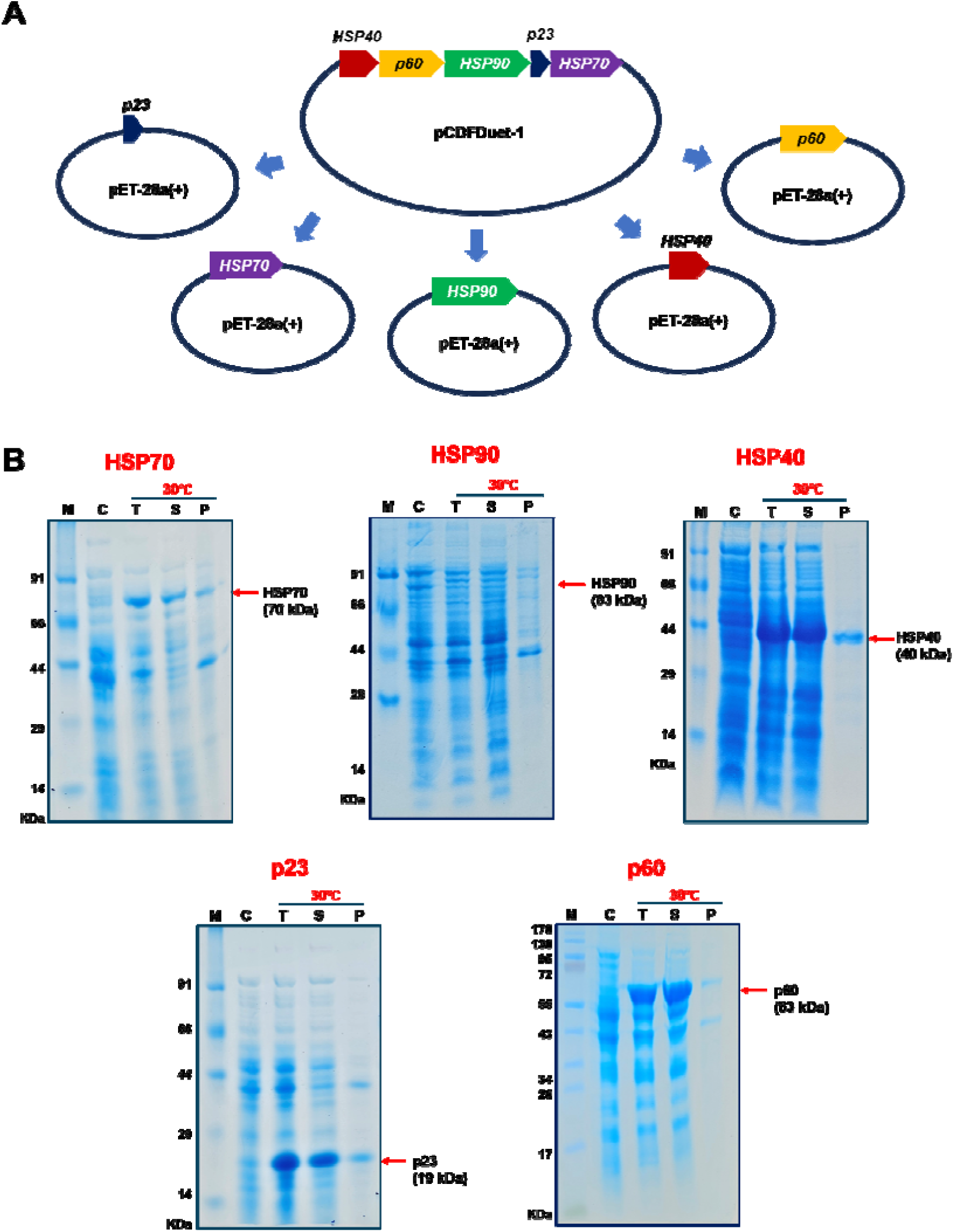
Subcloning and individual expression of HSP90 multichaperone complex components. **(A)** Schematic representation of the cloning strategy. Individual genes encoding the components of the HSP90 multichaperone complex (Hsp40, p60, HSP90, p23, and HSP70) were isolated from the polycistronic pCDFDuet-1 vector and independently cloned into separate pET-28a(+) expression vectors. **(B)** SDS-PAGE analysis demonstrating the independent expression of each recombinant protein. Protein expression for all constructs was induced at 30°C with 1 mM IPTG. For each gel, lanes represent: M, molecular weight marker; C, control (uninduced); T, total cell lysate; S, soluble fraction; P, pellet (insoluble fraction). Red arrows indicate the target proteins and their corresponding molecular weights: HSP70 (70 kDa), HSP90 (83 kDa), Hsp40 (40 kDa), p23 (19 kDa), and p60 (63 kDa).

**Figure S3.**
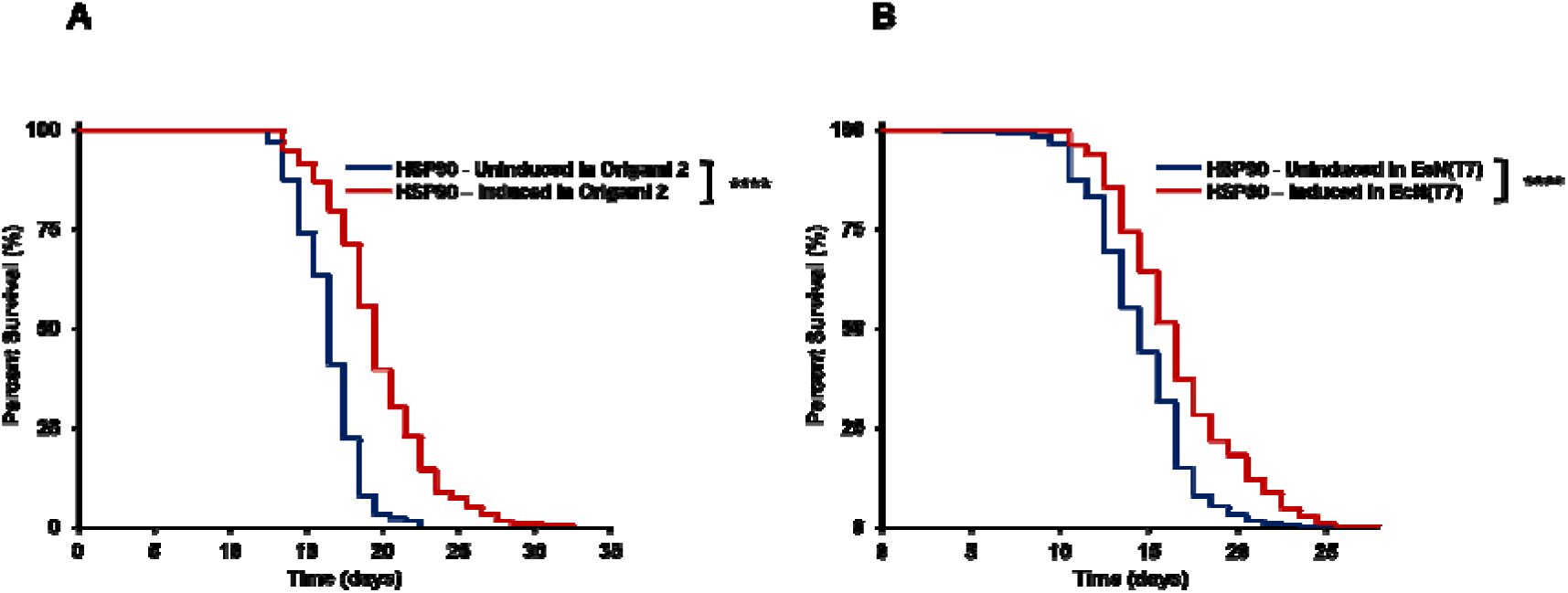
A diet of HSP90-expressing *E. coli* strains consistently extends wild-type *C. elegans*lifespan. For both panels, wild-type (N2) worms were fed bacteria harboring the HSP90 expression system, either induced to express HSP90 (1 mM IPTG at 30°C) or uninduced (lacking IPTG). **(A)** Kaplan-Meier survival curves demonstrate that the lifespan of N2 worms is significantly extended when fed HSP90-expressing *E. coli* Origami 2 (\*\*\*\**P* < 0.0001; uninduced *n* = 178, induced *n* = 178). **(B)** Similarly, Kaplan-Meier survival curves show a significantly extended lifespan for N2 worms fed HSP90-expressing *E. coli* Nissle 1917 [EcN(T7)] compared to uninduced controls (\*\*\*\**P* < 0.0001; uninduced *n* = 289, induced *n* = 324). (Statistical significance was determined using the Log-rank test. All lifespan assays represent three independent biological replicates).

### Supplementary Tables

**Table S1.** C. elegans strains utilized in this study.

| **Strain Designation** | **Genotype** |
|---|---|
| N2 | Wild-type (Bristol) |
| AM140 | *rmIs132 [unc-54p::Q35::YFP]* |
| CB1370 | *daf-2(e1370) III* |
| CF1038 | *daf-16(mu86) I* |
| DR412 | *daf-15(m81) unc-24(e138) IV* |
| KQ6 | *rict-1(mg360) II* |
| MQ130 | *clk-1(qm30) III* |
| MQ887 | *isp-1(qm150) IV* |
| QV225 | *skn-1(zj15) IV* |
| RB754 | *aak-2(ok524) X* |
| TJ1052 | *age-1(hx546) II* |
| TK22 | *mev-1(kn1) III* |
| VC475 | *hsp-16.2(gk249) V* |
| JT6130 | *hsp-90(p673) V* |
| PS3551 | *hsf-1(sy441) I* |

## References

1 Ohsawa, M., Uezono, Y. & Inui, A. Editorial: Ageing-Related Symptoms, Kampo Medicine, and Treatment. Front Nutr 8, 749320 (2021). 10.3389/fnut.2021.749320

2 López-Otín, C., Blasco, M. A., Partridge, L., Serrano, M. & Kroemer, G. Hallmarks of aging: An expanding universe. Cell 186, 243–278 (2023). 10.1016/j.cell.2022.11.001

3 Hipp, M. S., Kasturi, P. & Hartl, F. U. The proteostasis network and its decline in ageing. Nat Rev Mol Cell Biol 20, 421–435 (2019). 10.1038/s41580-019-0101-y

4 Labbadia, J. & Morimoto, R. I. The biology of proteostasis in aging and disease. Annu Rev Biochem 84, 435–464 (2015). 10.1146/annurev-biochem-060614-033955

5 Hartl, F. U. & Hayer-Hartl, M. Molecular chaperones in the cytosol: from nascent chain to folded protein. Science 295, 1852–1858 (2002). 10.1126/science.1068408

6 Hartl, F. U., Bracher, A. & Hayer-Hartl, M. Molecular chaperones in protein folding and proteostasis. Nature 475, 324–332 (2011). 10.1038/nature10317

7 Richter, K., Haslbeck, M. & Buchner, J. The heat shock response: life on the verge of death. Mol Cell 40, 253–266 (2010). 10.1016/j.molcel.2010.10.006

8 Chen, B., Retzlaff, M., Roos, T. & Frydman, J. Cellular strategies of protein quality control. Cold Spring Harb Perspect Biol 3, a004374 (2011). 10.1101/cshperspect.a004374

9 Höhfeld, J., Cyr, D. M. & Patterson, C. From the cradle to the grave: molecular chaperones that may choose between folding and degradation. EMBO Rep 2, 885–890 (2001). 10.1093/embo-reports/kve206

10 Finka, A. & Goloubinoff, P. Proteomic data from human cell cultures refine mechanisms of chaperone-mediated protein homeostasis. Cell Stress Chaperones 18, 591–605 (2013). 10.1007/s12192-013-0413-3

11 Taipale, M., Jarosz, D. F. & Lindquist, S. HSP90 at the hub of protein homeostasis: emerging mechanistic insights. Nat Rev Mol Cell Biol 11, 515–528 (2010). 10.1038/nrm2918

12 Bobkova, N. V. et al. Exogenous Hsp70 delays senescence and improves cognitive function in aging mice. Proc Natl Acad Sci U S A 112, 16006–16011 (2015). 10.1073/pnas.1516131112

13 Morán Luengo, T., Mayer, M. P. & Rüdiger, S. G. D. The Hsp70-Hsp90 Chaperone Cascade in Protein Folding. Trends Cell Biol 29, 164–177 (2019). 10.1016/j.tcb.2018.10.004

14 Mayer, M. P. & Bukau, B. Hsp70 chaperones: cellular functions and molecular mechanism. Cell Mol Life Sci 62, 670–684 (2005). 10.1007/s00018-004-4464-6

15 McClellan, A. J. et al. Diverse cellular functions of the Hsp90 molecular chaperone uncovered using systems approaches. Cell 131, 121–135 (2007). 10.1016/j.cell.2007.07.036

16 Schopf, F. H., Biebl, M. M. & Buchner, J. The HSP90 chaperone machinery. Nat Rev Mol Cell Biol 18, 345–360 (2017). 10.1038/nrm.2017.20

17 Backe, S. J., Sager, R. A., Woodford, M. R., Makedon, A. M. & Mollapour, M. Post-translational modifications of Hsp90 and translating the chaperone code. J Biol Chem 295, 11099–11117 (2020). 10.1074/jbc.REV120.011833

18 Han, S. Y. et al. Molecular Chaperone HSP90 Is Necessary to Prevent Cellular Senescence via Lysosomal Degradation of p14ARF. Cancer Res 77, 343–354 (2017). 10.1158/0008-5472.Can-16-0613

19 Meller, A. & Shalgi, R. The aging proteostasis decline: From nematode to human. Exp Cell Res 399, 112474 (2021). 10.1016/j.yexcr.2021.112474

20 Fuhrmann-Stroissnigg, H. et al. Identification of HSP90 inhibitors as a novel class of senolytics. Nat Commun 8, 422 (2017). 10.1038/s41467-017-00314-z

21 Hansen, D. T., Thiyagarajan, T., Larson, A. C. & Hansen, J. L. Telomerase repeat amplification protocol (TRAP) activity upon recombinant expression and purification of human telomerase in a bacterial system. Protein Expr Purif 123, 6–13 (2016). 10.1016/j.pep.2016.03.001

22 Fiege, K. & Frankenberg-Dinkel, N. Construction of a new T7 promoter compatible Escherichia coli Nissle 1917 strain for recombinant production of heme-dependent proteins. Microb Cell Fact 19, 190 (2020). 10.1186/s12934-020-01447-5

23 Nozaki, S. & Niki, H. Exonuclease III (XthA) Enforces In Vivo DNA Cloning of Escherichia coli To Create Cohesive Ends. J Bacteriol 201 (2019). 10.1128/jb.00660-18

24 Porta-de-la-Riva, M., Fontrodona, L., Villanueva, A. & Cerón, J. Basic Caenorhabditis elegans methods: synchronization and observation. J Vis Exp, e4019 (2012). 10.3791/4019

25 Han, S. K. et al. OASIS 2: online application for survival analysis 2 with features for the analysis of maximal lifespan and healthspan in aging research. Oncotarget 7, 56147–56152 (2016). 10.18632/oncotarget.11269

26 Pang, S. & Curran, S. P. Adaptive capacity to bacterial diet modulates aging in C. elegans. Cell Metab 19, 221–231 (2014). 10.1016/j.cmet.2013.12.005

27 Sánchez-Blanco, A. et al. Dietary and microbiome factors determine longevity in Caenorhabditis elegans. Aging (Albany NY) 8, 1513–1539 (2016). 10.18632/aging.101008

28 Bhat, A. et al. A diet of oxidative stress-adapted bacteria improves stress resistance and lifespan in C. elegans via p38-MAPK. Sci Adv 10, eadk8823 (2024). 10.1126/sciadv.adk8823

29 Padilla, C. J. et al. Profiling age-related muscle weakness and wasting: neuromuscular junction transmission as a driver of age-related physical decline. Geroscience 43, 1265–1281 (2021). 10.1007/s11357-021-00369-3

30 Borzuola, R. et al. Central and Peripheral Neuromuscular Adaptations to Ageing. J Clin Med 9 (2020). 10.3390/jcm9030741

31 Ben-Zvi, A., Miller, E. A. & Morimoto, R. I. Collapse of proteostasis represents an early molecular event in Caenorhabditis elegans aging. Proc Natl Acad Sci U S A 106, 14914–14919 (2009). 10.1073/pnas.0902882106

32 Walther, D. M. et al. Widespread Proteome Remodeling and Aggregation in Aging C. elegans. Cell 161, 919–932 (2015). 10.1016/j.cell.2015.03.032

33 Savola, E. et al. Testing evolutionary explanations for the lifespan benefit of dietary restriction in fruit flies (Drosophila melanogaster). Evolution 75, 450–463 (2021). 10.1111/evo.14146

34 Altintas, O., Park, S. & Lee, S. J. The role of insulin/IGF-1 signaling in the longevity of model invertebrates, C. elegans and D. melanogaster. BMB Rep 49, 81–92 (2016). 10.5483/bmbrep.2016.49.2.261

35 Murphy, C. T. & Hu, P. J. Insulin/insulin-like growth factor signaling in C. elegans. WormBook, 1–43 (2013). 10.1895/wormbook.1.164.1

36 Lin, K., Hsin, H., Libina, N. & Kenyon, C. Regulation of the Caenorhabditis elegans longevity protein DAF-16 by insulin/IGF-1 and germline signaling. Nat Genet 28, 139–145 (2001). 10.1038/88850

37 Morris, J. Z., Tissenbaum, H. A. & Ruvkun, G. A phosphatidylinositol-3-OH kinase family member regulating longevity and diapause in Caenorhabditis elegans. Nature 382, 536–539 (1996). 10.1038/382536a0

38 Weinkove, D., Halstead, J. R., Gems, D. & Divecha, N. Long-term starvation and ageing induce AGE-1/PI 3-kinase-dependent translocation of DAF-16/FOXO to the cytoplasm. BMC Biol 4, 1 (2006). 10.1186/1741-7007-4-1

39 Hsu, A. L., Murphy, C. T. & Kenyon, C. Regulation of aging and age-related disease by DAF-16 and heat-shock factor. Science 300, 1142–1145 (2003). 10.1126/science.1083701

40 Son, H. G. et al. Prefoldin 6 mediates longevity response from heat shock factor 1 to FOXO in C. elegans. Genes Dev 32, 1562–1575 (2018). 10.1101/gad.317362.118

41 Kaushik, N. et al. Transcriptome Analysis of Insulin Signaling-Associated Transcription Factors in C. elegans Reveal Their Genome-Wide Target Genes Specificity and Complexity. Int J Mol Sci 22 (2021). 10.3390/ijms222212462

42 Singh, V. & Aballay, A. Heat shock and genetic activation of HSF-1 enhance immunity to bacteria. Cell Cycle 5, 2443–2446 (2006). 10.4161/cc.5.21.3434

43 Barna, J. et al. Heat shock factor-1 intertwines insulin/IGF-1, TGF-β and cGMP signaling to control development and aging. BMC Dev Biol 12, 32 (2012). 10.1186/1471-213x-12-32

44 Rea, S. L., Wu, D., Cypser, J. R., Vaupel, J. W. & Johnson, T. E. A stress-sensitive reporter predicts longevity in isogenic populations of Caenorhabditis elegans. Nat Genet 37, 894–898 (2005). 10.1038/ng1608

45 Birnby, D. A. et al. A transmembrane guanylyl cyclase (DAF-11) and Hsp90 (DAF-21) regulate a common set of chemosensory behaviors in caenorhabditis elegans. Genetics 155, 85–104 (2000). 10.1093/genetics/155.1.85

46 Gaiser, A. M., Kaiser, C. J., Haslbeck, V. & Richter, K. Downregulation of the Hsp90 system causes defects in muscle cells of Caenorhabditis elegans. PLoS One 6, e25485 (2011). 10.1371/journal.pone.0025485

47 Greer, E. L. et al. An AMPK-FOXO pathway mediates longevity induced by a novel method of dietary restriction in C. elegans. Curr Biol 17, 1646–1656 (2007). 10.1016/j.cub.2007.08.047

48 Saxton, R. A. & Sabatini, D. M. mTOR Signaling in Growth, Metabolism, and Disease. Cell 168, 960–976 (2017). 10.1016/j.cell.2017.02.004

49 Johnson, S. C., Rabinovitch, P. S. & Kaeberlein, M. mTOR is a key modulator of ageing and age-related disease. Nature 493, 338–345 (2013). 10.1038/nature11861

50 Saxton, R. A. & Sabatini, D. M. mTOR Signaling in Growth, Metabolism, and Disease. Cell 169, 361–371 (2017). 10.1016/j.cell.2017.03.035

51 Soukas, A. A., Kane, E. A., Carr, C. E., Melo, J. A. & Ruvkun, G. Rictor/TORC2 regulates fat metabolism, feeding, growth, and life span in Caenorhabditis elegans. Genes Dev 23, 496–511 (2009). 10.1101/gad.1775409

52 An, J. H. & Blackwell, T. K. SKN-1 links C. elegans mesendodermal specification to a conserved oxidative stress response. Genes Dev 17, 1882–1893 (2003). 10.1101/gad.1107803

53 Blackwell, T. K., Steinbaugh, M. J., Hourihan, J. M., Ewald, C. Y. & Isik, M. SKN-1/Nrf, stress responses, and aging in Caenorhabditis elegans. Free Radic Biol Med 88, 290–301 (2015). 10.1016/j.freeradbiomed.2015.06.008

54 Yang, W. & Hekimi, S. A mitochondrial superoxide signal triggers increased longevity in Caenorhabditis elegans. PLoS Biol 8, e1000556 (2010). 10.1371/journal.pbio.1000556

55 Ishii, N. et al. A mutation in succinate dehydrogenase cytochrome b causes oxidative stress and ageing in nematodes. Nature 394, 694–697 (1998). 10.1038/29331

56 Ewbank, J. J. et al. Structural and functional conservation of the Caenorhabditis elegans timing gene clk-1. Science 275, 980–983 (1997). 10.1126/science.275.5302.980

57 Feng, J., Bussière, F. & Hekimi, S. Mitochondrial electron transport is a key determinant of life span in Caenorhabditis elegans. Dev Cell 1, 633–644 (2001). 10.1016/s1534-5807(01)00071-5

58 Oliveira, P. H. & Mairhofer, J. Marker-free plasmids for biotechnological applications - implications and perspectives. Trends Biotechnol 31, 539–547 (2013). 10.1016/j.tibtech.2013.06.001

59 Chen, J. et al. A survey of drug resistance bla genes originating from synthetic plasmid vectors in six Chinese rivers. Environ Sci Technol 46, 13448–13454 (2012). 10.1021/es302760s

60 Cheng, Y. Y. et al. Efficient plasmid transfer via natural competence in a microbial co-culture. Mol Syst Biol 19, e11406 (2023). 10.15252/msb.202211406

61 Amrofell, M. B. et al. Engineering E. coli strains using antibiotic-resistance-gene-free plasmids. Cell Rep Methods 3, 100669 (2023). 10.1016/j.crmeth.2023.100669

62 Kaushik, S. & Cuervo, A. M. Proteostasis and aging. Nat Med 21, 1406–1415 (2015). 10.1038/nm.4001

63 Yue, L. et al. Genetic analysis of viable Hsp90 alleles reveals a critical role in Drosophila spermatogenesis. Genetics 151, 1065–1079 (1999). 10.1093/genetics/151.3.1065

64 Fuhrmann-Stroissnigg, H., Niedernhofer, L. J. & Robbins, P. D. Hsp90 inhibitors as senolytic drugs to extend healthy aging. Cell Cycle 17, 1048–1055 (2018). 10.1080/15384101.2018.1475828

65 Brehme, M. et al. A chaperome subnetwork safeguards proteostasis in aging and neurodegenerative disease. Cell Rep 9, 1135–1150 (2014). 10.1016/j.celrep.2014.09.042

66 Kazemi-Esfarjani, P. & Benzer, S. Genetic suppression of polyglutamine toxicity in Drosophila. Science 287, 1837–1840 (2000). 10.1126/science.287.5459.1837

67 Chan, H. Y., Warrick, J. M., Gray-Board, G. L., Paulson, H. L. & Bonini, N. M. Mechanisms of chaperone suppression of polyglutamine disease: selectivity, synergy and modulation of protein solubility in Drosophila. Hum Mol Genet 9, 2811–2820 (2000). 10.1093/hmg/9.19.2811

68 Song, H. O. et al. C. elegans STI-1, the homolog of Sti1/Hop, is involved in aging and stress response. J Mol Biol 390, 604–617 (2009). 10.1016/j.jmb.2009.05.035

69 Mamat, U. et al. Detoxifying Escherichia coli for endotoxin-free production of recombinant proteins. Microb Cell Fact 14, 57 (2015). 10.1186/s12934-015-0241-5

70 Jeong, H., Kim, H. J. & Lee, S. J. Complete Genome Sequence of Escherichia coli Strain BL21. Genome Announc 3 (2015). 10.1128/genomeA.00134-15

71 Bessette, P. H., Aslund, F., Beckwith, J. & Georgiou, G. Efficient folding of proteins with multiple disulfide bonds in the Escherichia coli cytoplasm. Proc Natl Acad Sci U S A 96, 13703–13708 (1999). 10.1073/pnas.96.24.13703

72 Chiosis, G., Digwal, C. S., Trepel, J. B. & Neckers, L. Structural and functional complexity of HSP90 in cellular homeostasis and disease. Nat Rev Mol Cell Biol 24, 797–815 (2023). 10.1038/s41580-023-00640-9

73 Cardoso, V. M., Paredes, S. A. H., Campani, G., Gonçalves, V. M. & Zangirolami, T. C. ClearColi as a platform for untagged pneumococcal surface protein A production: cultivation strategy, bioreactor culture, and purification. Appl Microbiol Biotechnol 106, 1011–1029 (2022). 10.1007/s00253-022-11758-9

74 Revelles, O. et al. The carbon storage regulator (Csr) system exerts a nutrient-specific control over central metabolism in Escherichia coli strain Nissle 1917. PLoS One 8, e66386 (2013). 10.1371/journal.pone.0066386

75 Bansal, A., Zhu, L. J., Yen, K. & Tissenbaum, H. A. Uncoupling lifespan and healthspan in Caenorhabditis elegans longevity mutants. Proc Natl Acad Sci U S A 112, E277–286 (2015). 10.1073/pnas.1412192112

76 van Oosten-Hawle, P., Porter, R. S. & Morimoto, R. I. Regulation of organismal proteostasis by transcellular chaperone signaling. Cell 153, 1366–1378 (2013). 10.1016/j.cell.2013.05.015

77 Zou, J., Guo, Y., Guettouche, T., Smith, D. F. & Voellmy, R. Repression of heat shock transcription factor HSF1 activation by HSP90 (HSP90 complex) that forms a stress-sensitive complex with HSF1. Cell 94, 471–480 (1998). 10.1016/s0092-8674(00)81588-3

78 Kovács, D. et al. Age-dependent heat shock hormesis to HSF-1 deficiency suggests a compensatory mechanism mediated by the unfolded protein response and innate immunity in young Caenorhabditis elegans. Aging Cell 23, e14246 (2024). 10.1111/acel.14246

79 Maklakov, A. A. & Chapman, T. Evolution of ageing as a tangle of trade-offs: energy versus function. Proc Biol Sci 286, 20191604 (2019). 10.1098/rspb.2019.1604

80 Flatt, T. Survival costs of reproduction in Drosophila. Exp Gerontol 46, 369–375 (2011). 10.1016/j.exger.2010.10.008

81 Jenkins, N. L., McColl, G. & Lithgow, G. J. Fitness cost of extended lifespan in Caenorhabditis elegans. Proc Biol Sci 271, 2523–2526 (2004). 10.1098/rspb.2004.2897

82 Mair, W. & Dillin, A. Aging and survival: the genetics of life span extension by dietary restriction. Annu Rev Biochem 77, 727–754 (2008). 10.1146/annurev.biochem.77.061206.171059

83 Inoue, T. et al. Cell cycle control by daf-21/Hsp90 at the first meiotic prophase/metaphase boundary during oogenesis in Caenorhabditis elegans. Dev Growth Differ 48, 25–32 (2006). 10.1111/j.1440-169X.2006.00841.x

84 Wang, Y., Xu, S., Liu, J., Zhang, Y. & Guo, T. L. Regulation of lead toxicity by heat shock protein 90 (daf-21) is affected by temperature in Caenorhabditis elegans. Ecotoxicol Environ Saf 104, 317–322 (2014). 10.1016/j.ecoenv.2014.03.016

85 Somogyvári, M., Gecse, E. & Sőti, C. DAF-21/Hsp90 is required for C. elegans longevity by ensuring DAF-16/FOXO isoform A function. Sci Rep 8, 12048 (2018). 10.1038/s41598-018-30592-6

86 Morley, J. F. & Morimoto, R. I. Regulation of longevity in Caenorhabditis elegans by heat shock factor and molecular chaperones. Molecular biology of the cell 15, 657–664 (2004). 10.1091/mbc.e03-07-0532

87 Murphy, C. T. et al. Genes that act downstream of DAF-16 to influence the lifespan of Caenorhabditis elegans. Nature 424, 277–283 (2003). 10.1038/nature01789

88 Henderson, S. T. & Johnson, T. E. daf-16 integrates developmental and environmental inputs to mediate aging in the nematode Caenorhabditis elegans. Curr Biol 11, 1975–1980 (2001). 10.1016/s0960-9822(01)00594-2

89 Taipale, M. et al. A quantitative chaperone interaction network reveals the architecture of cellular protein homeostasis pathways. Cell 158, 434–448 (2014). 10.1016/j.cell.2014.05.039

90 Oh, S. W. et al. JNK regulates lifespan in Caenorhabditis elegans by modulating nuclear translocation of forkhead transcription factor/DAF-16. Proc Natl Acad Sci U S A 102, 4494–4499 (2005). 10.1073/pnas.0500749102

91 Lehtinen, M. K. et al. A conserved MST-FOXO signaling pathway mediates oxidative-stress responses and extends life span. Cell 125, 987–1001 (2006). 10.1016/j.cell.2006.03.046

92 Sidera, K., Samiotaki, M., Yfanti, E., Panayotou, G. & Patsavoudi, E. Involvement of cell surface HSP90 in cell migration reveals a novel role in the developing nervous system. J Biol Chem 279, 45379–45388 (2004). 10.1074/jbc.M405486200

93 Sidera, K. & Patsavoudi, E. Extracellular HSP90: conquering the cell surface. Cell Cycle 7, 1564–1568 (2008). 10.4161/cc.7.11.6054

94 Bhatia, A., O’Brien, K., Chen, M., Woodley, D. T. & Li, W. Keratinocyte-Secreted Heat Shock Protein-90alpha: Leading Wound Reepithelialization and Closure. Adv Wound Care (New Rochelle) 5, 176–184 (2016). 10.1089/wound.2014.0620

95 Wong, D. S. & Jay, D. G. Emerging Roles of Extracellular Hsp90 in Cancer. Adv Cancer Res 129, 141–163 (2016). 10.1016/bs.acr.2016.01.001

96 Poggio, P., Sorge, M., Seclì, L. & Brancaccio, M. Extracellular HSP90 Machineries Build Tumor Microenvironment and Boost Cancer Progression. Front Cell Dev Biol 9, 735529 (2021). 10.3389/fcell.2021.735529

97 Hance, M. W., Nolan, K. D. & Isaacs, J. S. The double-edged sword: conserved functions of extracellular hsp90 in wound healing and cancer. Cancers (Basel) 6, 1065–1097 (2014). 10.3390/cancers6021065

98 Sarkar, A. A. & Zohn, I. E. Hectd1 regulates intracellular localization and secretion of Hsp90 to control cellular behavior of the cranial mesenchyme. J Cell Biol 196, 789–800 (2012). 10.1083/jcb.201105101

99 Woodley, D. T. et al. Participation of the lipoprotein receptor LRP1 in hypoxia-HSP90alpha autocrine signaling to promote keratinocyte migration. J Cell Sci 122, 1495–1498 (2009). 10.1242/jcs.047894

100 Seclì, L. et al. Targeting the Extracellular HSP90 Co-Chaperone Morgana Inhibits Cancer Cell Migration and Promotes Anticancer Immunity. Cancer Res 81, 4794–4807 (2021). 10.1158/0008-5472.Can-20-3150

101 Thuringer, D. et al. Transactivation of the epidermal growth factor receptor by heat shock protein 90 via Toll-like receptor 4 contributes to the migration of glioblastoma cells. J Biol Chem 286, 3418–3428 (2011). 10.1074/jbc.M110.154823

102 Belsches-Jablonski, A. P. et al. Src family kinases and HER2 interactions in human breast cancer cell growth and survival. Oncogene 20, 1465–1475 (2001). 10.1038/sj.onc.1204205

103 Qin, Z., DeFee, M., Isaacs, J. S. & Parsons, C. Extracellular Hsp90 serves as a co-factor for MAPK activation and latent viral gene expression during de novo infection by KSHV. Virology 403, 92–102 (2010). 10.1016/j.virol.2010.03.052

104 Yochem, J., Tuck, S., Greenwald, I. & Han, M. A gp330/megalin-related protein is required in the major epidermis of Caenorhabditis elegans for completion of molting. Development 126, 597–606 (1999). 10.1242/dev.126.3.597

105 Pujol, N. et al. A reverse genetic analysis of components of the Toll signaling pathway in Caenorhabditis elegans. Curr Biol 11, 809–821 (2001). 10.1016/s0960-9822(01)00241-x

106 Davis, R. J. Signal transduction by the JNK group of MAP kinases. Cell 103, 239–252 (2000). 10.1016/s0092-8674(00)00116-1

107 Mukhopadhyay, A., Oh, S. W. & Tissenbaum, H. A. Worming pathways to and from DAF-16/FOXO. Exp Gerontol 41, 928–934 (2006). 10.1016/j.exger.2006.05.020

108 Dayalan Naidu, S., et al. Heat Shock Factor 1 Is a Substrate for p38 Mitogen-Activated Protein Kinases. Mol Cell Biol 36, 2403–2417 (2016). 10.1128/mcb.00292-16

109 Barna, J., Csermely, P. & Vellai, T. Roles of heat shock factor 1 beyond the heat shock response. Cell Mol Life Sci 75, 2897–2916 (2018). 10.1007/s00018-018-2836-6

110 Zhang, L. et al. Hsp90 interacts with AMPK and mediates acetyl-CoA carboxylase phosphorylation. Cell Signal 24, 859–865 (2012). 10.1016/j.cellsig.2011.12.001

111 Ohji, G. et al. Suppression of the mTOR-raptor signaling pathway by the inhibitor of heat shock protein 90 geldanamycin. J Biochem 139, 129–135 (2006). 10.1093/jb/mvj008

112 Delgoffe, G. M., Kole, T. P., Cotter, R. J. & Powell, J. D. Enhanced interaction between Hsp90 and raptor regulates mTOR signaling upon T cell activation. Mol Immunol 46, 2694–2698 (2009). 10.1016/j.molimm.2009.05.185

113 Jones, K. T., Greer, E. R., Pearce, D. & Ashrafi, K. Rictor/TORC2 regulates Caenorhabditis elegans fat storage, body size, and development through sgk-1. PLoS Biol 7, e60 (2009). 10.1371/journal.pbio.1000060

114 Wang, M. et al. The Therapeutic Roles of Recombinant Hsp90α on Cornea Epithelial Injury. Invest Ophthalmol Vis Sci 63, 30 (2022). 10.1167/iovs.63.2.30

115 Sarbassov, D. D., Guertin, D. A., Ali, S. M. & Sabatini, D. M. Phosphorylation and regulation of Akt/PKB by the rictor-mTOR complex. Science 307, 1098–1101 (2005). 10.1126/science.1106148

116 Xiao, R. et al. A genetic program promotes C. elegans longevity at cold temperatures via a thermosensitive TRP channel. Cell 152, 806–817 (2013). 10.1016/j.cell.2013.01.020

117 Mizunuma, M., Neumann-Haefelin, E., Moroz, N., Li, Y. & Blackwell, T. K. mTORC2-SGK-1 acts in two environmentally responsive pathways with opposing effects on longevity. Aging Cell 13, 869–878 (2014). 10.1111/acel.12248

118 Inoue, H. et al. The C. elegans p38 MAPK pathway regulates nuclear localization of the transcription factor SKN-1 in oxidative stress response. Genes Dev 19, 2278–2283 (2005). 10.1101/gad.1324805

119 Schieber, M. & Chandel, N. S. ROS function in redox signaling and oxidative stress. Curr Biol 24, R453–462 (2014). 10.1016/j.cub.2014.03.034

120 Ristow, M. & Schmeisser, K. Mitohormesis: Promoting Health and Lifespan by Increased Levels of Reactive Oxygen Species (ROS). Dose Response 12, 288–341 (2014). 10.2203/dose-response.13-035.Ristow

121 Zanphorlin, L. M. et al. Heat Shock Protein 90 kDa (Hsp90) Has a Second Functional Interaction Site with the Mitochondrial Import Receptor Tom70. J Biol Chem 291, 18620–18631 (2016). 10.1074/jbc.M115.710137

122 Budas, G. R., Churchill, E. N., Disatnik, M. H., Sun, L. & Mochly-Rosen, D. Mitochondrial import of PKCepsilon is mediated by HSP90: a role in cardioprotection from ischaemia and reperfusion injury. Cardiovasc Res 88, 83–92 (2010). 10.1093/cvr/cvq154

123 Felkai, S. et al. CLK-1 controls respiration, behavior and aging in the nematode Caenorhabditis elegans. Embo j 18, 1783–1792 (1999). 10.1093/emboj/18.7.1783

124 Branicky, R., Nguyen, P. A. & Hekimi, S. Uncoupling the pleiotropic phenotypes of clk-1 with tRNA missense suppressors in Caenorhabditis elegans. Mol Cell Biol 26, 3976–3985 (2006). 10.1128/mcb.26.10.3976-3985.2006

125 Senoo-Matsuda, N. et al. A defect in the cytochrome b large subunit in complex II causes both superoxide anion overproduction and abnormal energy metabolism in Caenorhabditis elegans. J Biol Chem 276, 41553–41558 (2001). 10.1074/jbc.M104718200

126 Kenyon, C., Chang, J., Gensch, E., Rudner, A. & Tabtiang, R. A C. elegans mutant that lives twice as long as wild type. Nature 366, 461–464 (1993). 10.1038/366461a0

127 Kimura, K. D., Tissenbaum, H. A., Liu, Y. & Ruvkun, G. daf-2, an insulin receptor-like gene that regulates longevity and diapause in Caenorhabditis elegans. Science 277, 942–946 (1997). 10.1126/science.277.5328.942

128 López-Perrote, A., Serna, M. & Llorca, O. Maturation and Assembly of mTOR Complexes by the HSP90-R2TP-TTT Chaperone System: Molecular Insights and Mechanisms. Subcell Biochem 104, 459–483 (2024). 10.1007/978-3-031-58843-3_17

129 Nieto-Miguel, T., Gajate, C., González-Camacho, F. & Mollinedo, F. Proapoptotic role of Hsp90 by its interaction with c-Jun N-terminal kinase in lipid rafts in edelfosine-mediated antileukemic therapy. Oncogene 27, 1779–1787 (2008). 10.1038/sj.onc.1210816

130 Yang, R., Kang, Y., Duan, J., Zou, C. & Wu, Q. The p38 MAPK/PMK-1 Pathway Is Required for Resistance to Nocardia farcinica Infection in Caenorhabditis elegance. Pathogens 11 (2022). 10.3390/pathogens11101071

131 Gopal, U. et al. A novel extracellular Hsp90 mediated co-receptor function for LRP1 regulates EphA2 dependent glioblastoma cell invasion. PLoS One 6, e17649 (2011). 10.1371/journal.pone.0017649

132 Tsen, F. et al. Extracellular heat shock protein 90 signals through subdomain II and the NPVY motif of LRP-1 receptor to Akt1 and Akt2: a circuit essential for promoting skin cell migration in vitro and wound healing in vivo. Mol Cell Biol 33, 4947–4959 (2013). 10.1128/mcb.00559-13

133 Pearce, L. R., Komander, D. & Alessi, D. R. The nuts and bolts of AGC protein kinases. Nat Rev Mol Cell Biol 11, 9–22 (2010). 10.1038/nrm2822

134 Mora, A., Komander, D., van Aalten, D. M. & Alessi, D. R. PDK1, the master regulator of AGC kinase signal transduction. Semin Cell Dev Biol 15, 161–170 (2004). 10.1016/j.semcdb.2003.12.022

135 Alessi, D. R. et al. Characterization of a 3-phosphoinositide-dependent protein kinase which phosphorylates and activates protein kinase Balpha. Curr Biol 7, 261–269 (1997). 10.1016/s0960-9822(06)00122-9

136 Park, J. et al. Serum and glucocorticoid-inducible kinase (SGK) is a target of the PI 3-kinase-stimulated signaling pathway. Embo j 18, 3024–3033 (1999). 10.1093/emboj/18.11.3024

137 García-Martínez, J. M. & Alessi, D. R. mTOR complex 2 (mTORC2) controls hydrophobic motif phosphorylation and activation of serum- and glucocorticoid-induced protein kinase 1 (SGK1). Biochem J 416, 375–385 (2008). 10.1042/bj20081668

138 Hertweck, M., Göbel, C. & Baumeister, R. C. elegans SGK-1 is the critical component in the Akt/PKB kinase complex to control stress response and life span. Dev Cell 6, 577–588 (2004). 10.1016/s1534-5807(04)00095-4

139 Tullet, J. M. et al. Direct inhibition of the longevity-promoting factor SKN-1 by insulin-like signaling in C. elegans. Cell 132, 1025–1038 (2008). 10.1016/j.cell.2008.01.030

140 Chiang, W. C., Ching, T. T., Lee, H. C., Mousigian, C. & Hsu, A. L. HSF-1 regulators DDL-1/2 link insulin-like signaling to heat-shock responses and modulation of longevity. Cell 148, 322–334 (2012). 10.1016/j.cell.2011.12.019

141 Brunet, A. et al. Protein kinase SGK mediates survival signals by phosphorylating the forkhead transcription factor FKHRL1 (FOXO3a). Mol Cell Biol 21, 952–965 (2001). 10.1128/mcb.21.3.952-965.2001

142 Kobayashi, T. & Cohen, P. Activation of serum- and glucocorticoid-regulated protein kinase by agonists that activate phosphatidylinositide 3-kinase is mediated by 3-phosphoinositide-dependent protein kinase-1 (PDK1) and PDK2. Biochem J 339 (Pt 2), 319–328 (1999).

143 Tullet, J. M. A. et al. The SKN-1/Nrf2 transcription factor can protect against oxidative stress and increase lifespan in C. elegans by distinct mechanisms. Aging Cell 16, 1191–1194 (2017). 10.1111/acel.12627

144 Wei, H. et al. Heat shock protein 90: biological functions, diseases, and therapeutic targets. MedComm (2020) 5, e470 (2024). 10.1002/mco2.470

145 Tariq, M., Nussbaumer, U., Chen, Y., Beisel, C. & Paro, R. Trithorax requires Hsp90 for maintenance of active chromatin at sites of gene expression. Proc Natl Acad Sci U S A 106, 1157–1162 (2009). 10.1073/pnas.0809669106

146 Chen, B. & Wagner, A. Hsp90 is important for fecundity, longevity, and buffering of cryptic deleterious variation in wild fly populations. BMC Evol Biol 12, 25 (2012). 10.1186/1471-2148-12-25

147 Whitesell, L. & Lindquist, S. L. HSP90 and the chaperoning of cancer. Nat Rev Cancer 5, 761–772 (2005). 10.1038/nrc1716

148 Kamal, A. et al. A high-affinity conformation of Hsp90 confers tumour selectivity on Hsp90 inhibitors. Nature 425, 407–410 (2003). 10.1038/nature01913

149 Janssens, G. E. et al. Transcriptomics-Based Screening Identifies Pharmacological Inhibition of Hsp90 as a Means to Defer Aging. Cell Rep 27, 467–480.e466 (2019). 10.1016/j.celrep.2019.03.044

150 Prodromou, C. et al. Identification and structural characterization of the ATP/ADP-binding site in the Hsp90 molecular chaperone. Cell 90, 65–75 (1997). 10.1016/s0092-8674(00)80314-1

151 Pearl, L. H., Prodromou, C. & Workman, P. The Hsp90 molecular chaperone: an open and shut case for treatment. Biochem J 410, 439–453 (2008). 10.1042/bj20071640

152 Neckers, L. & Workman, P. Hsp90 molecular chaperone inhibitors: are we there yet? Clin Cancer Res 18, 64–76 (2012). 10.1158/1078-0432.Ccr-11-1000

153 Solit, D. B. & Chiosis, G. Development and application of Hsp90 inhibitors. Drug Discov Today 13, 38–43 (2008). 10.1016/j.drudis.2007.10.007

154 Corsi, A. K., Wightman, B. & Chalfie, M. A Transparent window into biology: A primer on Caenorhabditis elegans. WormBook, 1–31 (2015). 10.1895/wormbook.1.177.1

155 Naranjo-Galindo, F. J., Ai, R., Fang, E. F., Nilsen, H. L. & SenGupta, T. C. elegans as an Animal Model to Study the Intersection of DNA Repair, Aging and Neurodegeneration. Front Aging 3, 916118 (2022). 10.3389/fragi.2022.916118

156 Somogyvári, M., Khatatneh, S. & Sőti, C. Hsp90: From Cellular to Organismal Proteostasis. Cells 11 (2022). 10.3390/cells11162479

157 Garcia-Toscano, L. et al. Decreased Hsp90 activity protects against TDP-43 neurotoxicity in a C. elegans model of amyotrophic lateral sclerosis. PLoS Genet 20, e1011518 (2024). 10.1371/journal.pgen.1011518

158 Charbonneau, M. R., Isabella, V. M., Li, N. & Kurtz, C. B. Developing a new class of engineered live bacterial therapeutics to treat human diseases. Nat Commun 11, 1738 (2020). 10.1038/s41467-020-15508-1

159 Stecher, B. et al. Gut inflammation can boost horizontal gene transfer between pathogenic and commensal Enterobacteriaceae. Proc Natl Acad Sci U S A 109, 1269–1274 (2012). 10.1073/pnas.1113246109

160 von Wintersdorff, C. J. et al. Dissemination of Antimicrobial Resistance in Microbial Ecosystems through Horizontal Gene Transfer. Front Microbiol 7, 173 (2016). 10.3389/fmicb.2016.00173

161 Lynch, J. P., Goers, L. & Lesser, C. F. Emerging strategies for engineering Escherichia coli Nissle 1917-based therapeutics. Trends Pharmacol Sci 43, 772–786 (2022). 10.1016/j.tips.2022.02.002

162 Kim, J. & Moon, Y. Worm-Based Alternate Assessment of Probiotic Intervention against Gut Barrier Infection. Nutrients 11 (2019). 10.3390/nu11092146

163 Helmy, Y. A., Kassem, II, Kumar, A. & Rajashekara, G. In Vitro Evaluation of the Impact of the Probiotic E. coli Nissle 1917 on Campylobacter jejuni’s Invasion and Intracellular Survival in Human Colonic Cells. Front Microbiol 8, 1588 (2017). 10.3389/fmicb.2017.01588

164 Dubois, C. et al. Exposure of Caenorhabditis elegans to Dietary Nε-Carboxymethyllysine Emphasizes Endocytosis as a New Route for Intestinal Absorption of Advanced Glycation End Products. Nutrients 13 (2021). 10.3390/nu13124398

165 Sato, K., Norris, A., Sato, M. & Grant, B. D. C. elegans as a model for membrane traffic. WormBook, 1–47 (2014). 10.1895/wormbook.1.77.2

166 Chen, C. C. et al. RAB-10 is required for endocytic recycling in the Caenorhabditis elegans intestine. Mol Biol Cell 17, 1286–1297 (2006). 10.1091/mbc.e05-08-0787

167 Grant, B. & Hirsh, D. Receptor-mediated endocytosis in the Caenorhabditis elegans oocyte. Mol Biol Cell 10, 4311–4326 (1999). 10.1091/mbc.10.12.4311

168 Wang, J. et al. C. elegans ciliated sensory neurons release extracellular vesicles that function in animal communication. Curr Biol 24, 519–525 (2014). 10.1016/j.cub.2014.01.002

169 Winston, W. M., Sutherlin, M., Wright, A. J., Feinberg, E. H. & Hunter, C. P. Caenorhabditis elegans SID-2 is required for environmental RNA interference. Proc Natl Acad Sci U S A 104, 10565–10570 (2007). 10.1073/pnas.0611282104

170 Hodge, F., Bajuszova, V. & van Oosten-Hawle, P. The Intestine as a Lifespan- and Proteostasis-Promoting Signaling Tissue. Front Aging 3, 897741 (2022). 10.3389/fragi.2022.897741

171 Nono, M. et al. Intestine-to-Germline Transmission of Epigenetic Information Intergenerationally Ensures Systemic Stress Resistance in C. elegans. Cell Rep 30, 3207–3217.e3204 (2020). 10.1016/j.celrep.2020.02.050

172 Veuthey, T. et al. The neurohormone tyramine stimulates the secretion of an insulin-like peptide from the Caenorhabditis elegans intestine to modulate the systemic stress response. PLoS Biol 23, e3002997 (2025). 10.1371/journal.pbio.3002997

173 Zhang, P., Judy, M., Lee, S. J. & Kenyon, C. Direct and indirect gene regulation by a life-extending FOXO protein in C. elegans: roles for GATA factors and lipid gene regulators. Cell Metab 17, 85–100 (2013). 10.1016/j.cmet.2012.12.013

